# Challenging the right-hemisphere assumption in post-stroke pragmatics: largely comparable impairment profiles across lesion sides

**DOI:** 10.64898/2026.08.06.742887

**Authors:** Simone Gastaldon, Fortunata Romeo, Chiara Barattieri di San Pietro, Natalia Chumakova, Daniela D’Imperio, Sara Lago, Sara Nordio, Ilaria Parrotta, Marco Rigoni, Valentina Bambini, Giorgio Arcara

**Affiliations:** Brain Rhythms and Cognition Group, BCBL – Basque Center on Cognition, Brain and Language, Donostia-San Sebastián, Spain; Laboratory of Neurolinguistics and Experimental Pragmatics (NEPLab), University School for Advanced Studies IUSS, Pavia, Italy; Study Center of Neurocognitive Rehabilitation “Villa Miari”, Santorso, Vicenza, Italy; Department of Philosophy, Sociology, Education and Applied Psychology (FISPPA), University of Padua, Padua, Italy; IRCCS San Camillo Hospital, Venice, Italy; Department of General Psychology (DPG), University of Padua, Padua, Italy

**Keywords:** pragmatics, non-literal language, stroke, aphasia, lesion lateralization

## Abstract

Traditional views assume that pragmatic deficits after stroke, which compromise the interpretation of communicative intentions and non-literal meanings, follow damage to the right hemisphere (RHD), with left-hemisphere damage (LHD) primarily linked to aphasia and structural language impairment. To examine hemispheric contributions to post-stroke pragmatic profiles, we assessed 99 stroke patients (40 LHD, including 14 with aphasia of minimal-to-moderate severity; 59 RHD) and 60 healthy controls with the Assessment of Pragmatic Abilities and Cognitive Substrates (APACS). While stroke patients overall performed worse than controls, LHD and RHD profiles were largely comparable across three converging analyses: (1) permutation tests revealed no hemispheric differences except on the two tasks requiring expressive components (Interview and Figurative Language 2), which in turn lowered the composites (APACS Production and Total); (2) equivalence testing established equivalence for most measures, with only these same tasks and composites remaining inconclusive; and (3) unsupervised clustering did not group patients by lesion side. Theory of Mind was robustly associated with pragmatic performance in both groups, whereas structural language abilities related specifically to LHD performance and general cognition only to RHD. Excluding aphasic LHD patients strengthened the evidence for comparable profiles, indicating that aphasic LHD patients largely drove the residual differences. In conclusion, primary pragmatic impairment, especially in the receptive domain, emerged comparably after LHD and RHD, with the only residual LHD disadvantage limited to tasks demanding open verbal output. These findings challenge the assumption of right-hemispheric specialization for pragmatics, stressing the need for pragmatic assessment in all post-stroke patients.

## Introduction

Stroke is a leading cause of acquired motor and cognitive disability (Herpich & Rincon, 2020; Karamyan, 2023), frequently leaving lingering impairments across motor, sensory, cognitive, and psychological domains that require sustained rehabilitation. It is commonly accompanied by communication deficits, with substantial consequences for social participation, quality of life, and daily activities (Bays, 2001; Fridriksson & Hillis, 2021; Rousseaux et al., 2010). The most documented post-stroke communication disorder is aphasia, arising primarily from left-hemisphere damage (LHD) to regions supporting phonology, lexical semantics, and syntax (Sheppard and Sebastian, 2021). However, communication difficulties in stroke extend well beyond the structural language deficits that define aphasia, entering the domain of pragmatics.

Pragmatics refers to the ability to integrate linguistic content with contextual information and infer interlocutors’ intentions in communication (Grice, 1989; Sperber & Wilson, 1995). It includes organizing coherent and informative discourse, managing conversational exchange, and understanding figurative, non-literal and indirect forms of language such as metaphors, idioms, humor, and irony (Angeleri et al., 2012; Arcara & Bambini, 2016; Bischetti, Frau, & Bambini, 2024; Frau et al., 2025). When these abilities are impaired, as often follows brain damage, the consequences can be severe, resulting in loss of referential links, literal interpretation of figurative expressions, and violations of conversational rules (Deighton et al., 2020; Cummings, 2021).

### Left vs. Right Hemisphere Debate in Pragmatics: An Overview

The history of clinical pragmatics is closely tied to the study of right-hemisphere damage (RHD). Before the 1960s, right-hemisphere lesions were thought to leave language intact, since the classical aphasia syndromes were clearly left-lateralized; the diffusion of pragmatic theory, initially articulated by Grice (1989) and Searle (1979), then gave a vocabulary to a clinical observation that already existed: RHD patients struggled not with the form of language but with its use in context (García, Ferré, & Joanette, 2021). Through the 1970s and 1980s the right hemisphere came to be seen as the locus of pragmatic ability, partly because clinicians needed to separate these difficulties from classical LHD aphasia (Gardner & Brownell, 1986; Joanette et al., 1990; Blake, 2017), with early reports of impaired metaphor (Winner & Gardner, 1977), humor and irony (Brownell et al., 1983), and narrative and conversational organization (Joanette et al., 1990). Out of this work grew the “right-hemisphere hypothesis” of pragmatics, which dominated the field for almost three decades.

That picture is too simple even for RHD itself. Pragmatic impairments are neither necessary nor uniform consequences of right-hemisphere stroke. Although they are common, as about half of RHD patients show one or more communication difficulties, rising to roughly 78% in acute settings (Blake et al., 2026; García, Ferré, & Joanette, 2021; Parola et al., 2016), their manifestations vary considerably, from pervasive deficits to mild prosodic or conversational problems, and some RHD patients show no detectable disorder (Ferré et al., 2012).

Pragmatic impairment is also not specific to RHD. Over the past two decades it has been documented across many neurological, neurodegenerative, and psychiatric conditions: traumatic brain injury (Rowley et al., 2017; Arcara et al., 2020), schizophrenia (Champagne-Lavau, Stip, & Joanette, 2007; Bambini et al., 2016b; Bambini et al., 2020), multiple sclerosis (Carotenuto et al., 2018; Lago et al., 2022; Nordio et al., 2025), amyotrophic lateral sclerosis (Bambini et al., 2016a), Parkinson’s disease (Montemurro et al., 2019), and adult dyslexia (Cappelli et al., 2018), among others (Cummings, 2021; Bischetti et al., 2024). Its prevalence differs from one condition to the next but is rarely absent (Bischetti et al., 2024; Frau et al., 2025). In a cross-diagnostic study of 454 participants across seven clinical groups, Frau et al. (2025) found receptive pragmatic skills impaired in every group relative to controls, and expressive pragmatic skills impaired in the four groups of neurological origin (Parkinson’s disease, right-hemisphere stroke, amyotrophic lateral sclerosis, and traumatic brain injury). Pragmatic difficulty, thus, qualifies more as a broad feature of brain disorder, rather than a signature of RHD.

The left hemisphere tells a similar story. Pragmatic difficulties can accompany aphasia in LHD or occur without it: Borod et al. (2000) found LHD patients more impaired than RHD patients in the pragmatic appropriateness of discourse, and Kasher et al. (1999) reported clear LHD deficits relative to controls. Direct comparisons give a mixed result that depends on the pragmatic component and the task. Using gestural (extralinguistic) tasks, Cutica, Bucciarelli, and Bara (2006) found pragmatic performance better preserved after left-than after right-hemisphere damage, whereas Sidtis and Yang (2017), studying formulaic language, saw right-hemisphere involvement only in speech output elicitation and not in multiple-choice formats. Others found no differences (Spaccavento et al., 2024). Recovery of structural language after LHD does not guarantee recovery of pragmatic communication (Coelho & Flewellyn, 2003), which indicates that pragmatics relies in part on resources outside the classical language network. Neural evidence points the same way: non-literal language is supported jointly by the linguistic and the Theory of Mind (ToM; Premack and Woodruff, 1978) systems (Bischetti, Frau, & Bambini, 2024; Paunov et al., 2022; Tomasello et al., 2025), with a proposed pragmatic white-matter pathway along the arcuate fasciculus (Catani & Bambini, 2014), and yet the two systems do not fully overlap (Forbes Schieche et al., 2025). Seen in this light, the early dominance of the right hemisphere looks partly methodological: aphasic LHD patients were commonly excluded, which inflated the apparent RHD effect (Zaidel et al., 2002; Ferstl, 2008).

The rationale for the exclusion of LHD patients reflects an important clinical distinction. Pragmatic disorders can be primary or secondary: a secondary disorder is one in which pragmatic skills suffer as a consequence of structural language or general cognitive deficits, whereas a primary disorder does not arise from such deficits (Cummings, 2021). This distinction matters most for LHD. When aphasia is present, some pragmatic failures may be secondary to structural damage rather than signs of genuine pragmatic impairment, since grammatical or lexico-semantic problems can confound figurative-language testing and non-fluent aphasia can impede production tasks (Cummings, 2021). Because primary and secondary impairment are hard to tell apart, aphasic LHD patients have largely been left out of clinical pragmatics research. Exclusion, however, does not settle the matter: pragmatic difficulty in aphasic LHD can exceed what structural impairment alone would predict (Glosser & Goodglass, 1990; Fridriksson et al., 2006). Non-aphasic LHD patients are also underrepresented, since they fit neither the aphasia rehabilitation pathway nor the more clinically salient RHD profile (García, Ferré, & Joanette, 2021). Whether pragmatic impairment in LHD is an independent disruption, a secondary consequence, or both therefore remains open.

Even where pragmatic impairment is established, the cognitive mechanisms behind it remain debated, because pragmatic competence does not operate in isolation: context-based communication draws on a distributed set of resources (Mar, 2004). Core linguistic abilities provide the material on which inference operates, which matters in particular for LHD patients, whose linguistic deficits may constrain the expression of otherwise preserved pragmatic capacities (Arcara & Bambini, 2016; Bischetti, Frau, & Bambini, 2024). Executive functions help select contextually appropriate meanings and integrate information across a discourse, and executive dysfunction has itself been proposed as a source of pragmatic impairment (Champagne-Lavau & Joanette, 2009; Martin & McDonald, 2003; Cummings, 2021; Tsolakopoulos et al., 2023). Abstract reasoning supports figurative language comprehension, where meaning must be carried from one conceptual domain to another (Chiappe & Chiappe, 2007). ToM has drawn the most attention, since recovering a speaker’s intended meaning in irony, indirect requests, or non-literal language requires representing their mental states (Sperber & Wilson, 2002), and ToM-based accounts have been especially prominent for autism, schizophrenia, and RHD (Cummings, 2015; Champagne-Lavau & Joanette, 2009). Its role, however, is partial and variable. Frau et al. (2025) found the ToM-comprehension association moderate and significant only in some groups, with no association between ToM and pragmatic production, and recent syntheses describe the pragmatics-ToM link as flexible rather than fixed, varying across pragmatic phenomena, populations, and tasks (Bambini & Lecce, 2025). By contrast, Papafragou and Grigoroglou (2025) see no reason to abandon the assumption that mentalizing supports pragmatic interpretation, attributing apparent dissociations to task demands and to the gap between having ToM and deploying it in a given context. Despite these differences, one practical point is recurring: pragmatics and its candidate substrates are related but distinct, and should be measured separately (Bosco, Tirassa, & Gabbatore, 2018; Bischetti et al., 2024).

This complexity has a direct consequence for lesion studies. If pragmatic processing draws on several systems, which are differently distributed at the neural level (Catani & Bambini, 2014; Bischetti, Frau, & Bambini, 2024), then the cognitive profile supporting pragmatic performance should itself depend on lesion side, with preserved linguistic resources weighing more heavily after LHD and ToM, executive, or reasoning components after RHD. Whether LHD and RHD pragmatic profiles are comparable is therefore only half the question; the other half is whether similar pragmatic performance rests on different cognitive foundations in the two groups. Addressing both requires a standardized instrument that measures pragmatics, accompanied by measures of the cognitive domains that contribute to pragmatic performance. We therefore adopted the Assessment of Pragmatic Abilities and Cognitive Substrates (APACS; Arcara & Bambini, 2016), a validated battery which assesses discourse production and figurative-language comprehension in Italian-speaking adults and has documented sensitivity across diverse clinical populations (Frau et al., 2025). The use of APACS is recommended in combination with cognitive assessment: accordingly, we complemented the pragmatic battery with standardized measures of the principal domains underlying pragmatics, including Theory of Mind, linguistic abilities, abstract reasoning, and executive functions.

### The present study

If post-stroke pragmatic impairment is in fact bilateral, attributing it to right-hemisphere damage alone risks misdirecting rehabilitation and leaving deficits unrecognized, above all in LHD patients, whose pragmatic abilities have been overshadowed by the clinical focus on aphasia. Clarifying the role of lesion side therefore matters both for rehabilitation and for how pragmatic deficits are understood across neurological conditions, and, more broadly, it helps dispel the view of the brain as a set of isolated functional modules rather than a highly interconnected system (Pessoa, 2022).

The present study examines pragmatic abilities in stroke patients with left- or right-hemisphere damage (LHD and RHD), assessed with the APACS. Unlike most prior work, the sample is not restricted to non-aphasic patients: to separate structural-language impairment from genuine pragmatic difficulty, every analysis is run in parallel on the full sample and on the subsample excluding aphasic patients. Given the evidence for bilateral pragmatic networks and the underestimated role of the left hemisphere, we expected pragmatic impairment in both groups, with possible differences in its magnitude or profile.

Accordingly, the present study pursues three aims: (1) To characterize the commonalities and differences in the pragmatic profiles of LHD and RHD patients, addressing aspects of communication too often neglected in LHD; (2) To test whether pragmatic abilities relate to other cognitive domains differently depending on the lesioned hemisphere, clarifying which cognitive components support pragmatic competence and whether their relative weight varies between LHD and RHD; (3) To establish whether pragmatic impairment in LHD survives the exclusion of aphasic cases, revealing potential primary pragmatic deficits in a clinically neglected population. Exploratorily, we also assessed patient’s functional communication via the Communication Outcome After Stroke questionnaire (Bambini et al., 2017), to investigate the relationship between perceived communicative effectiveness and actual pragmatic ability.

## Materials and Methods

### Participants

Participants were Italian-speaking stroke patients recruited at two Italian centers: IRCCS Ospedale San Camillo (Lido, Venice) and Istituzione Comunale Villa Miari (Santorso, Vicenza). Eligibility required being a native Italian speaker aged 18 or older, having sustained a single unilateral stroke confirmed by medical records, being in the post-acute stage (at least 3 months post-stroke), and being able to provide written informed consent. Exclusion criteria were severe impairment on the Denomination or Comprehension subtests of the Aachen Aphasia Test (AAT), any disability preventing participation, and a history of psychiatric or neurological disorders unrelated to the stroke. The research was performed in accordance with the principles stated in the Declaration of Helsinki and was approved by the Ethical Committee of Area Sud-Ovest Veneto (Protocol Code: PLR - Prog. 330CET).

A total of 99 stroke survivors participated in the study (age: M = 61.85, SD = 12.79; education: M = 11.07 years, SD = 4.15; 36 women), 40 with LHD (age: M = 61.20, SD = 11.56; education: M = 11.44, SD = 4.35; 18 women) and 59 with RHD (age: M = 62.29, SD = 13.64; education: M = 10.83, SD = 4.03; 18 women). The two groups did not differ in age (t = −0.43, p = 0.67), education (t = 0.69, p = 0.49), or sex distribution (χ² = 1.58, p = 0.208). Within the LHD group, 14 patients were categorized as aphasic, with Aachen Aphasia Test (AAT) performance indicating generally moderate-to-minimal residual structural language impairment and a single severe score, in Repetition (mean ± SD: Token Test 15.2 ± 12.1 errors; Repetition 125.5 ± 24.3; Writing 68.4 ± 20.5; Naming 103.9 ± 13.4; Comprehension 99.9 ± 11.1). Because all analyses were run in parallel on an aphasia-excluded sample, we also report its demographics: with the 14 aphasic patients removed, the LHD group (n = 26; age: M = 61.58, SD = 12.13; education: M = 12.28, SD = 4.16; 15 women) still matched the RHD group in age (t = −0.24, p = 0.812) and education (t = 1.47, p = 0.148), although the sex distribution became unbalanced (58% vs. 31% women; χ² = 4.53, p = 0.033). Figure 1 summarizes relevant demographic, clinical and anatomical data for the stroke sample.

**Figure 1.**
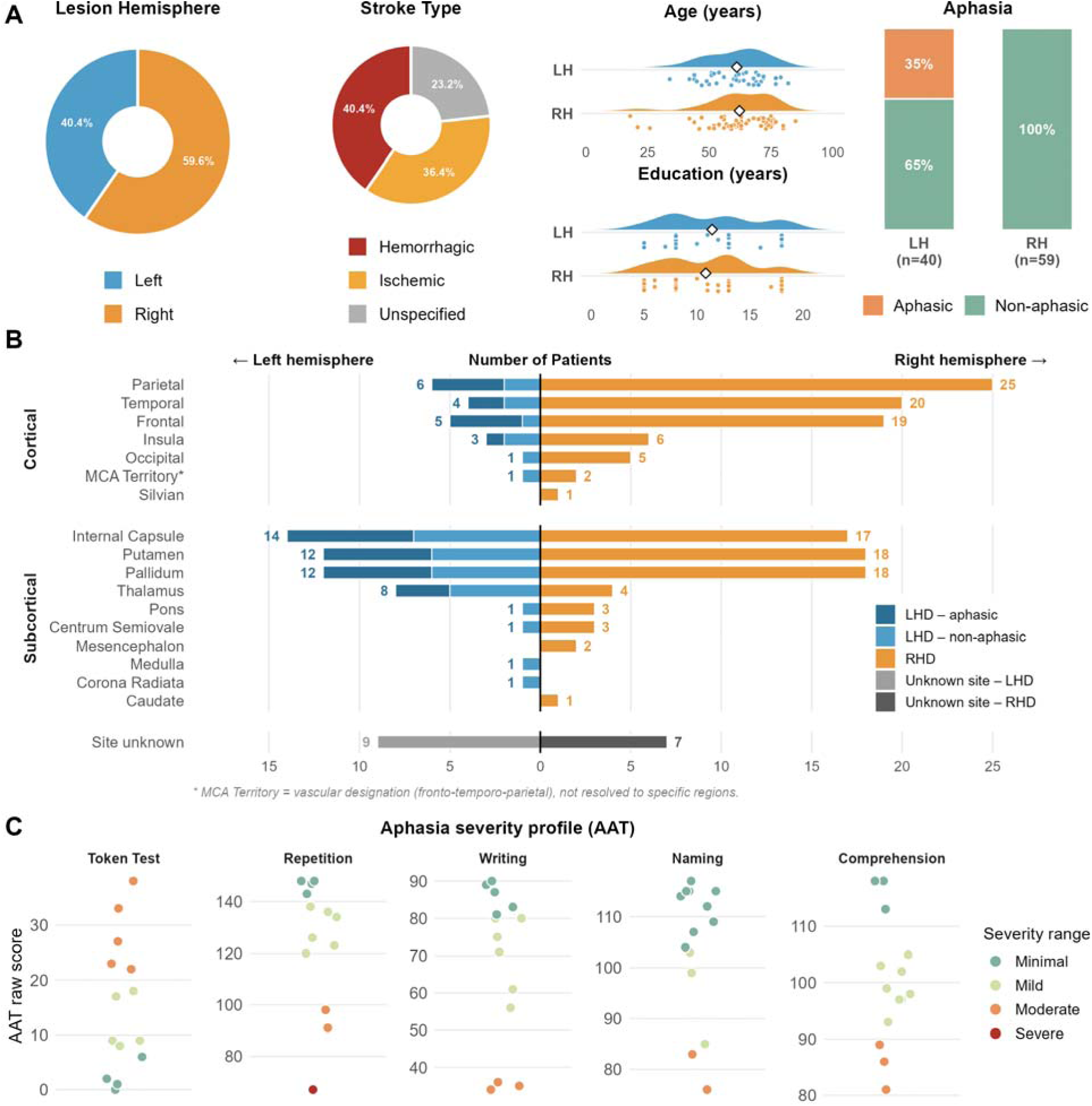
Sample characteristics and lesion site description of stroke patients (N = 99; LHD = 40, RHD = 59). **(A)** Demographic and clinical variables. Donut charts: lesion hemisphere and stroke type. Raincloud plots: age and education (years) by hemisphere group (LH, blue; RH, orange), where the half violin is the density, each dot one patient, and the open diamond the group mean. Stacked bars: proportion of aphasic and non-aphasic patients within each group. **(B)** Butterfly plot of lesion frequency across brain regions for LHD (blue) and RHD (orange), grouped into cortical and subcortical structures and sorted by total frequency within each group. The left branch is partitioned by aphasia status (dark blue = aphasic LHD, light blue = non-aphasic LHD). Bars count region mentions rather than patients, so a multi-region lesion contributes to more than one row. The bottom row (grey) shows patients whose lesion site was not documented (9 LHD, 7 RHD); lesion data were available for 83 of 99 patients, with anatomical nomenclature standardized from the medical records by one author (FR). *MCA (middle cerebral artery) Territory is a vascular designation (fronto-temporo-parietal) not resolved to specific regions. **(C)** AAT performance for the 14 aphasic patients (all LHD), one point per patient, subtests on independent y-axes (raw-score ranges differ), colored by severity band. The Token Test is an error score (higher = worse); the other four increase with performance.

A control group of 60 healthy participants was also recruited (age: M = 61.67, SD = 11.07; education: M = 11.25 years, SD = 4.40; 36 women). Controls and stroke patients were comparable in age (t = 0.10, p = 0.925) and education (t = −0.25, p = 0.801), whereas women were overrepresented among controls (60% vs. 36%; χ² = 7.50, p = 0.006). Previous normative data indicate that APACS performance is not affected by sex (Arcara & Bambini, 2016), suggesting that this imbalance is unlikely to have materially influenced the comparisons.

### Assessment materials

Pragmatic abilities were assessed with the Assessment of Pragmatic Abilities and Cognitive Substrates (APACS; Arcara & Bambini, 2016), which evaluates the two primary pragmatic domains, discourse and non-literal language, in Italian. It is organized into production and comprehension sections comprising six tasks (Interview, Description, Narratives, Figurative Language 1, Humor, and Figurative Language 2), from which three composite scores are derived (Production, Comprehension, Total; see Figure 2).

**Figure 2.**
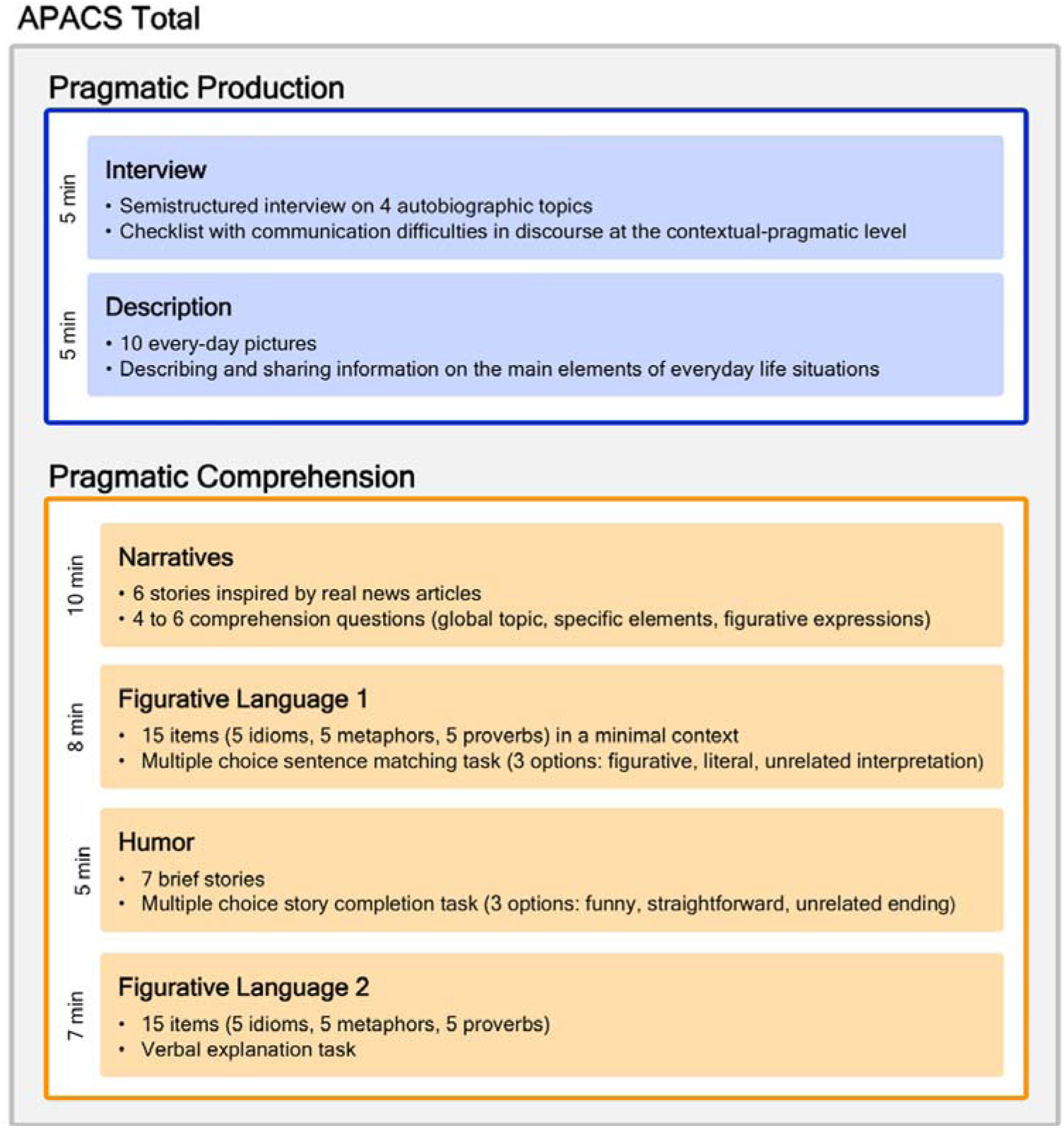
Structure of the APACS test: a Production section (blue, two tasks) and a Comprehension section (orange, four tasks). Interview: semi-structured conversation, scored for verbal pragmatics (speech, informativeness, information flow), paralinguistic dimensions, and grammar/vocabulary errors (max 44). Description: salient elements of ten everyday photographs (max 48). Narratives: discourse and narrative comprehension via open, yes/no, and non-literal explanation questions (max 56). Figurative Language 1: non-literal inference through multiple-choice explanations (max 15). Humor: verbal humor comprehension, multiple-choice (max 7). Figurative Language 2: verbal explanation of 15 figurative items, scored 0-2 each (max 30). Figure taken from Arcara and Bambini, *Frontiers in Psychology*, 2016, licensed under CC BY (DOI: <u>10.3389/fpsyg.2016.00070</u>).

A subset of patients was also administered tests assessing domains closely related to pragmatics. This subsampling reflected practical constraints typical of clinical stroke research, including assessment burden, fatigue, and limited clinical availability (Campbell et al., 2015). Analyses involving these measures were therefore restricted to participants with available data, with sample sizes reported for each analysis. We assessed Theory of Mind (ToM; Frank, 2018) with the Story-Based Empathy Task (SET; Dodich et al., 2015); structural linguistic skills with the Language subtest of the Addenbrooke’s Cognitive Examination – Revised (ACE-R Language; Mioshi et al., 2006); and abstract reasoning and executive functions with Raven’s Colored Progressive Matrices (Raven 47; Carlesimo et al., 1996; Roca et al., 2012). These were selected to tap the cognitive domains most relevant to pragmatic processing, following standard neuropsychological assessment practice (Lezak et al., 2012; Mondini, Cappelletti, & Arcara, 2022). The patient’s perceived functional communication effectiveness and its impact on quality of life was also assessed (Communication Outcome After Stroke, COAST; Bambini et al., 2017).

### Statistical analyses

All analyses were performed in R (R Core Team, 2023).

#### Permutation tests

We first compared patient performance on each APACS subtest and composite with healthy controls using non-parametric permutation testing (20,000 resamples of the mean difference), robust to the skewed distributions of these measures. Then we applied the same procedure to the LHD vs RHD comparison. Each measure was analyzed on an available-case basis. Effect sizes were quantified using Cliff’s δ, with 95% confidence intervals from 5,000 bootstrap resamples (Cliff, 1993; Meissel & Yao, 2024). P-values are reported uncorrected and corrected across the nine measures using the False Discovery Rate procedure (FDR; Benjamini & Hochberg, 1995). All comparisons were run in parallel on the full sample and on a subsample excluding aphasic patients.

#### Equivalence tests

Because a non-significant difference test cannot establish comparability (it indicates only a failure to detect a difference) we complemented the permutation tests with equivalence testing using the two one-sided tests procedure (TOST; Lakens, 2017; Lakens, Scheel, & Isager, 2018) to ask whether the differences are small enough to be practically negligible. Equivalence testing has only two outcomes, “equivalent” or “inconclusive”, and the latter is never interpreted as evidence of a difference. For each measure the TOST evaluates H_01_: μ_LHD_ − μ_RHD_ ≤ −ε and H_02_: μ_LHD_ − μ_RHD_ ≥ +ε; rejecting both at α = 0.05 places the mean difference within [−ε, +ε], equivalent to saying that its 90% CI lies entirely within ±ε. Given unequal group sizes, tests used the Welch (Satterthwaite) formulation, computed with the *tsum_TOST* function of the TOSTER package (Caldwell, 2022; Lakens, 2017). Rather than fixing ε as an arbitrary fraction of the maximum score, we anchored it to each measure’s standard error of measurement (SEM), estimated from the reliable-change tables of the APACS normative study (Supplementary Tables 2.1–2.9 in Arcara & Bambini, 2016; Crawford & Garthwaite, 2006): since each threshold is the product of the prediction error and z = 1.645, dividing the mean absolute distance between a tabulated score and its threshold by z recovers the SEM (non-computable cells excluded). Analyses used available cases and were run in parallel on the full and aphasia-excluded samples.

#### Patients’ profiling

For a clinical description, each individual score was classified as below or above the clinical cut-off for every measure, using the demographically adjusted cut-offs of Arcara and Bambini (2016) (Supplementary Tables 3.1-3.9 in Arcara & Bambini, 2016), selected accordingly to each patient’s age and education (Description being the only demographically invariant cut-off, as no demographic effect was found for it in the normative study). These cut-offs, derived with the Crawford and Garthwaite (2006) method, indicate less than 5% probability of observing that performance in the healthy normative sample. Based on these classifications, we computed the sample prevalence of below-threshold performance overall and by lesion side, summarizing the results in a prevalence bar plot (Figure 3B). Individual profiles were shown in a raster plot ordered by lesion side and number of impaired measures (Supplementary Figure 3). Profiling was conducted on an available-case basis: the group-level bar plot was generated both including and excluding aphasic patients, while aphasic patients were explicitly flagged in the individual-level profiling.

**Figure 3.**
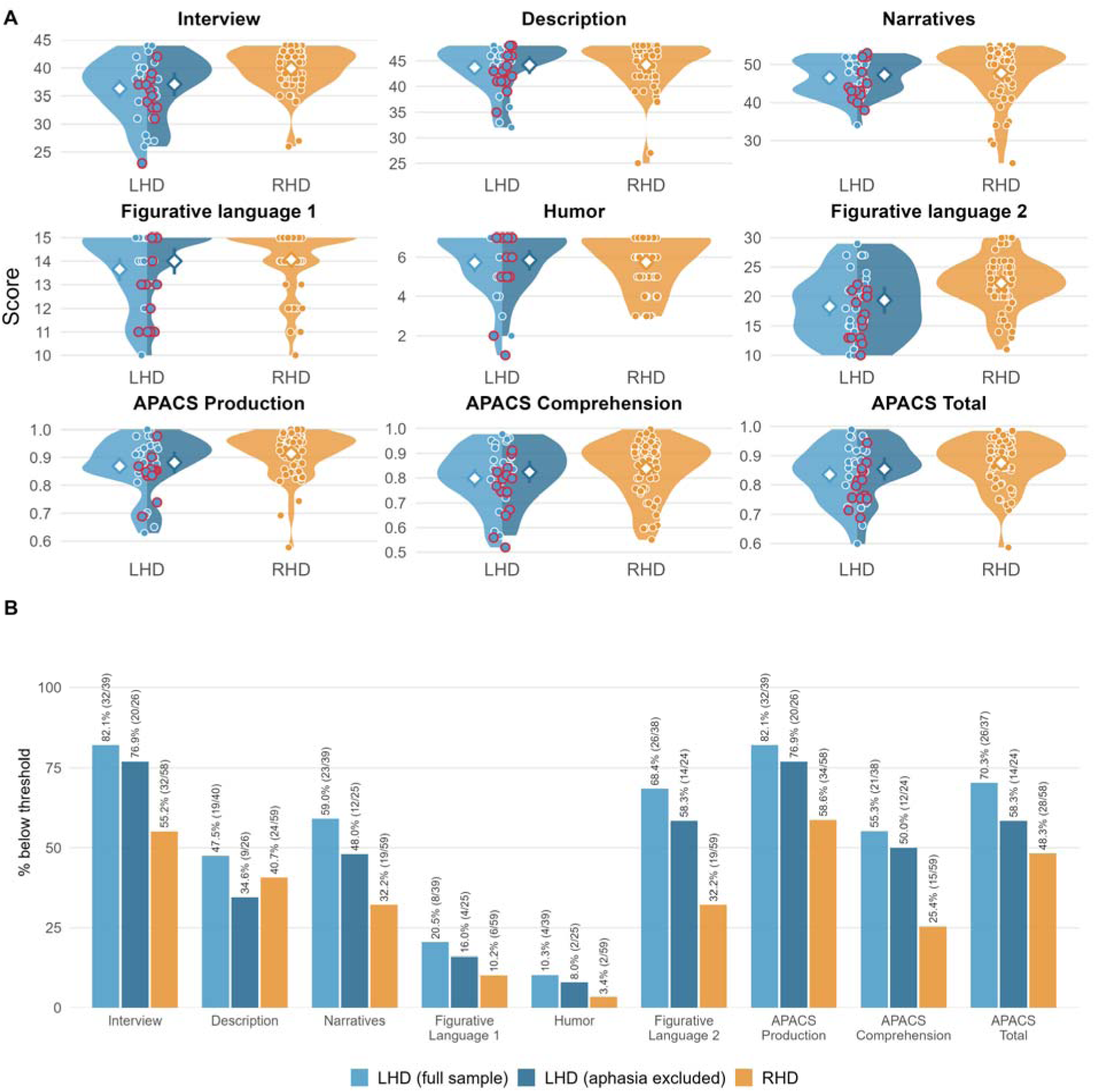
APACS performance and deficit prevalence in LHD and RHD patient samples. **(A)** APACS score distributions across the nine measures. Individual scores are dots; the group mean with its 95% CI (mean ± 1.96·SE) is a white diamond with a vertical bar; aphasic LHD patients are outlined in red. The LHD violin is split into two halves, colored as in panel B: light blue = whole LHD group, dark blue = LHD after excluding aphasic patients, each with its own mean. RHD is a single full violin (orange). **(B)** Prevalence of below-threshold performance, as grouped bars giving, for each measure, the percentage scoring below the published normative cut-off (5th percentile; Arcara & Bambini, 2016) in the full LHD sample (light blue), the aphasia-excluded LHD sample (dark blue), and RHD (orange; shown once, since no aphasic patient had right-hemisphere damage). Labels give the percentage and, in parentheses, the number below threshold over the number assessed. Denominators vary across measures and groups because percentages are computed on available cases.

#### K-means clustering

To further explore similarities and differences in pragmatic skills between LHD and RHD patients, we examined whether unsupervised clustering in the APACS scores recovers the lesion-side grouping. If patients with lesions in the same hemisphere can be grouped on the basis of pragmatic performance alone, this would support differential effects of lateralization; a failure to recover lesion side would point to comparable outcomes. APACS task scores were z-standardized and a Euclidean distance matrix computed. We applied k-means clustering with k = 2 (50 random initializations), assessing the match to lesion side via best-match accuracy, the φ coefficient and a χ² test, with average silhouette width indexing cohesion. Stability was evaluated across 200 re-runs. The procedure was run on the full sample and repeated excluding aphasic patients.

#### Correlations and robustness diagnostics

We examined associations between pragmatic performance and the additional cognitive assessments. Associations between APACS Total and the cognitive measures (SET, ACE-R Language, Raven) were estimated separately for LHD and RHD using Pearson’s correlation on an available-case basis, so pairwise sample sizes differ across associations. Within each group, the three p-values were FDR-adjusted (Benjamini & Hochberg, 1995), with significance at FDR < 0.05. To evaluate stability beyond simple statistical significance, we applied complementary robustness diagnostics. First, Spearman’s correlation was computed alongside Pearson’s, with an absolute discrepancy below 0.10 indicating agreement. Second, non-parametric bootstrap 95% CIs (1000 resamples) were derived, with bootstrap-stability defined as the interval excluding zero. Third, leave-one-out (LOO) sensitivity was assessed by recomputing each correlation omitting one observation; LOO-stability required SD below 0.10 with all estimates retaining sign. Fourth, the influence of observations was quantified using Cook’s distance, flagging values exceeding the 4/n threshold. These four criteria were combined into a composite robustness score (0-4), classified as high (≥3), moderate (2), or low (≤1). The same procedure was applied to the whole-sample APACS Total-COAST correlation; as a single test, its significance was evaluated against an uncorrected 0.05 threshold.

## Results

### APACS in stroke vs controls

As expected, stroke survivors performed significantly worse than controls on most APACS measures, the exceptions being Humor and Figurative Language 1 (see full results in Supplementary Table 1 and Supplementary Figures 1 and 2). The differences found in seven out of nine measures were reliable and robust, all surviving FDR correction (p_FDR_ < 0.001) with Cliff’s δ ranging from medium to large (−0.42 to −0.80). The largest deficits fell on production- and discourse-based measures (APACS Production, δ = −0.80; Interview, δ = −0.74; Figurative Language 2, δ = −0.59), whereas the two non-differing tasks, Figurative Language 1 (δ = −0.07, p_FDR_ = 0.286) and Humor (δ = −0.08, p_FDR_ = 0.423), are multiple-choice comprehension tasks on which both groups performed near ceiling.

### APACS in LHD vs RHD

#### Patients’ profiling

Below-threshold performance was common in both groups and followed a clear domain gradient, highest on discourse- and production-based measures (Interview, Narratives, Figurative Language 2, APACS Production) and the composites, lowest on Humor and Figurative Language 1 (Figure 3). In the full sample, prevalence was numerically higher in LHD than RHD on all nine measures, with the widest gaps on Interview, Figurative Language 2, and the APACS composites, and the narrowest where the two groups performed similarly (Description, Humor, Figurative Language 1). Excluding aphasic patients lowered deficit prevalence in the LHD sample on every measure but preserved the overall pattern of higher prevalence in LHD relative to RHD patients, with the exception of Description, where RHD performed slightly worse than LHD. Individual profiles are in Supplementary Figure 3.

#### Group differences

In the full sample, three measures survived FDR correction, all lower in LHD: Figurative Language 2 and Interview (both δ = −0.43) and APACS Production (δ = −0.30); APACS Total was significant before correction only, and the remaining five measures did not differ (all p_FDR_ ≥ 0.18; Table 1, Figure 4A,B). When aphasic patients were excluded, no measure survived correction (Table 1, Figure 4D,E): Interview and Figurative Language 2 remained the largest effects but attenuated to δ = −0.29 (significant uncorrected only), and the APACS Production and Total effects disappeared.

**Figure 4.**
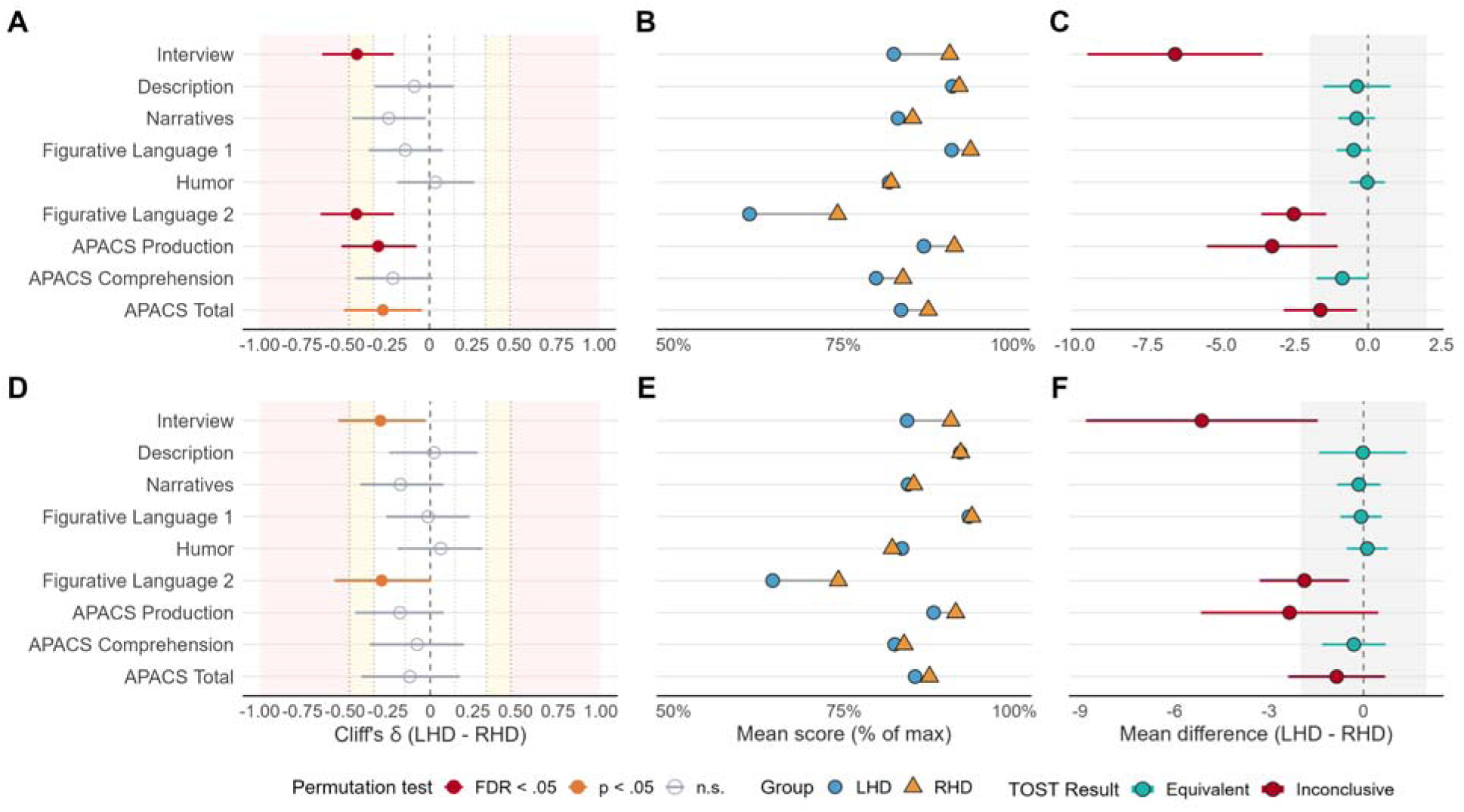
Permutation tests and equivalence tests of pragmatic performance in LHD versus RHD stroke patients across the nine APACS measures, in the full sample (A-C) and after excluding patients with aphasia (D-F). **(A, D)** Effect sizes (Cliff’s δ) for the LHD vs RHD contrast with 95% bootstrap CIs (5,000 resamples); negative = lower in LHD. Point color encodes the permutation-test outcome (20,000 resamples): red = significant after FDR correction, orange = significant before correction only, grey = non-significant. Shading marks effect-size benchmarks (Vargha & Delaney, 2000: medium |δ| ≥ .33; large |δ| ≥ .47), with dotted lines at ±.15, ±.33, ±.47. **(B, E)** Group means. Mean scores expressed as percentage of the maximum, for LHD (blue circles) and RHD (orange triangles) patients. **(C, F)** Equivalence testing (two one-sided tests, TOST). For each measure, the point is the LHD-RHD mean difference and the colored bar its 90% confidence interval; the grey band is the equivalence zone ±ε. The x-axis is expressed in units of each measure’s standard error of measurement (SEM): the mean difference and its 90% CI bounds are divided by that measure’s SEM. Teal = “equivalent” (90% CI entirely within the ±2 SEM band; both one-sided TOST *p* < 0.05); red = “inconclusive”, i.e. equivalence could not be established at this margin. Available cases: LHD *n* = 38-40 (25-26 excluding aphasia), RHD *n* = 58-59

**Table 1.**
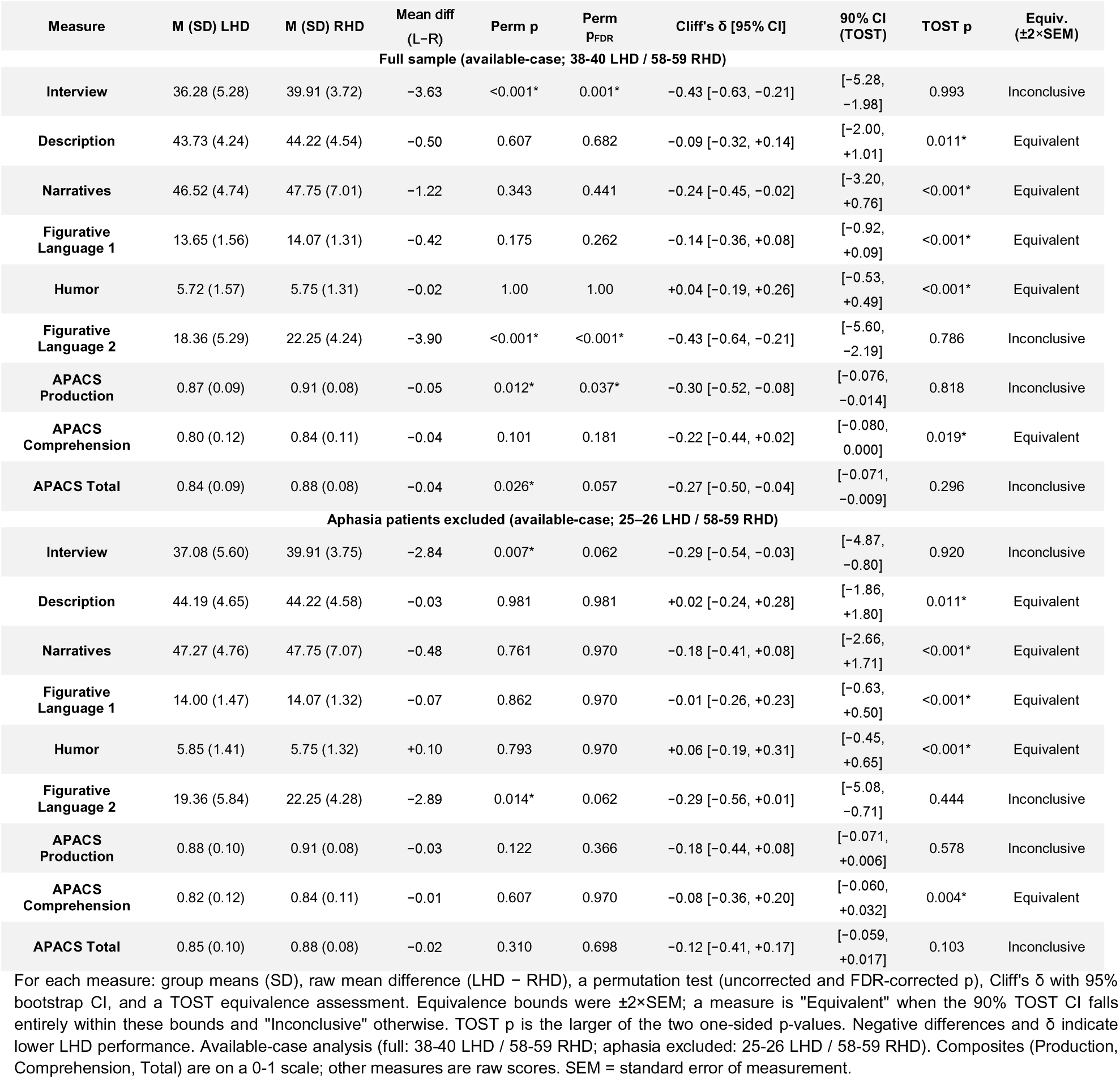
Comparisons (permutation and equivalence tests) of APACS performance in LHD and RHD patients, for the full sample and after excluding aphasic patients.

| Measure | M (SD) LHD | M (SD) RHD | Mean diff (L-R) | Perm p | Perm $p_{FDR}$ | Cliff's $\delta$ [95% CI] | 90% CI (TOST) | TOST p | Equiv. ($\pm 2 \times \text{SEM}$) |
| --- | --- | --- | --- | --- | --- | --- | --- | --- | --- |
| **Full sample (available-case; 38-40 LHD / 58-59 RHD)** |  |  |  |  |  |  |  |  |  |
| Interview | 36.28 (5.28) | 39.91 (3.72) | -3.63 | <0.001* | 0.001* | -0.43 [-0.63, -0.21] | [-5.28, -1.98] | 0.993 | Inconclusive |
| Description | 43.73 (4.24) | 44.22 (4.54) | -0.50 | 0.607 | 0.682 | -0.09 [-0.32, +0.14] | [-2.00, +1.01] | 0.011* | Equivalent |
| Narratives | 46.52 (4.74) | 47.75 (7.01) | -1.22 | 0.343 | 0.441 | -0.24 [-0.45, -0.02] | [-3.20, +0.76] | <0.001* | Equivalent |
| Figurative Language 1 | 13.65 (1.56) | 14.07 (1.31) | -0.42 | 0.175 | 0.262 | -0.14 [-0.36, +0.08] | [-0.92, +0.09] | <0.001* | Equivalent |
| Humor | 5.72 (1.57) | 5.75 (1.31) | -0.02 | 1.00 | 1.00 | +0.04 [-0.19, +0.26] | [-0.53, +0.49] | <0.001* | Equivalent |
| Figurative Language 2 | 18.36 (5.29) | 22.25 (4.24) | -3.90 | <0.001* | <0.001* | -0.43 [-0.64, -0.21] | [-5.60, -2.19] | 0.786 | Inconclusive |
| APACS Production | 0.87 (0.09) | 0.91 (0.08) | -0.05 | 0.012* | 0.037* | -0.30 [-0.52, -0.08] | [-0.076, -0.014] | 0.818 | Inconclusive |
| APACS Comprehension | 0.80 (0.12) | 0.84 (0.11) | -0.04 | 0.101 | 0.181 | -0.22 [-0.44, +0.02] | [-0.080, 0.000] | 0.019* | Equivalent |
| APACS Total | 0.84 (0.09) | 0.88 (0.08) | -0.04 | 0.026* | 0.057 | -0.27 [-0.50, -0.04] | [-0.071, -0.009] | 0.296 | Inconclusive |
| **Aphasia patients excluded (available-case; 25-26 LHD / 58-59 RHD)** |  |  |  |  |  |  |  |  |  |
| Interview | 37.08 (5.60) | 39.91 (3.75) | -2.84 | 0.007* | 0.062 | -0.29 [-0.54, -0.03] | [-4.87, -0.80] | 0.920 | Inconclusive |
| Description | 44.19 (4.65) | 44.22 (4.58) | -0.03 | 0.981 | 0.981 | +0.02 [-0.24, +0.28] | [-1.86, +1.80] | 0.011* | Equivalent |
| Narratives | 47.27 (4.76) | 47.75 (7.07) | -0.48 | 0.761 | 0.970 | -0.18 [-0.41, +0.08] | [-2.66, +1.71] | <0.001* | Equivalent |
| Figurative Language 1 | 14.00 (1.47) | 14.07 (1.32) | -0.07 | 0.862 | 0.970 | -0.01 [-0.26, +0.23] | [-0.63, +0.50] | <0.001* | Equivalent |
| Humor | 5.85 (1.41) | 5.75 (1.32) | +0.10 | 0.793 | 0.970 | +0.06 [-0.19, +0.31] | [-0.45, +0.65] | <0.001* | Equivalent |
| Figurative Language 2 | 19.36 (5.84) | 22.25 (4.28) | -2.89 | 0.014* | 0.062 | -0.29 [-0.56, +0.01] | [-5.08, -0.71] | 0.444 | Inconclusive |
| APACS Production | 0.88 (0.10) | 0.91 (0.08) | -0.03 | 0.122 | 0.366 | -0.18 [-0.44, +0.08] | [-0.071, +0.006] | 0.578 | Inconclusive |
| APACS Comprehension | 0.82 (0.12) | 0.84 (0.11) | -0.01 | 0.607 | 0.970 | -0.08 [-0.36, +0.20] | [-0.060, +0.032] | 0.004* | Equivalent |
| APACS Total | 0.85 (0.10) | 0.88 (0.08) | -0.02 | 0.310 | 0.698 | -0.12 [-0.41, +0.17] | [-0.059, +0.017] | 0.103 | Inconclusive |

For each measure: group means (SD), raw mean difference (LHD – RHD), a permutation test (uncorrected and FDR-corrected p), Cliff's $\delta$ with 95% bootstrap CI, and a TOST equivalence assessment. Equivalence bounds were $\pm 2 \times \text{SEM}$ ; a measure is "Equivalent" when the 90% TOST CI falls entirely within these bounds and "Inconclusive" otherwise. TOST p is the larger of the two one-sided p-values. Negative differences and $\delta$ indicate lower LHD performance. Available-case analysis (full: 38-40 LHD / 58-59 RHD; aphasia excluded: 25-26 LHD / 58-59 RHD). Composites (Production, Comprehension, Total) are on a 0-1 scale; other measures are raw scores. SEM = standard error of measurement.

#### Group equivalences

In the full sample, five measures were equivalent (Description, Narratives, Figurative Language 1, Humor, APACS Comprehension) and four inconclusive, each with a 90% CI lying entirely below zero: Interview, Figurative Language 2, APACS Production, and APACS Total (Figure 4C). Excluding aphasic patients differentiated this pattern. The APACS Production and Total intervals now crossed zero (Production [-0.071, +0.006]; Total [-0.059, +0.017]), so their inconclusive status reflected limited precision rather than a reliable difference, whereas Interview and Figurative Language 2 stayed below zero, marking a residual LHD disadvantage on the two open-speech tasks (Figure 4F). Overall, LHD and RHD patients were statistically equivalent on most pragmatic measures, with the only robust gaps on the open-speech tasks, which narrow once aphasia is removed.

#### Clustering

In both analyses the clusters separated by overall severity, not lesion side (Figure 5): one scored above the mean on all nine measures and the other below, with the widest gap on the composites. Crucially, each cluster contained a mix of LHD and RHD patients, failing to group by lesion side. Association with lesion side was weak and non-significant (full sample φ = 0.215, p = 0.059; aphasia excluded φ = 0.082, p = 0.636), with modest silhouette widths (0.41 and 0.43). Excluding aphasic patients, who were all LHD and did not form a separate group but were distributed across the lower-performing cluster, weakened the lateralization further. The unsupervised structure is thus organized by global pragmatic impairment rather than lateralization, consistent with a bilateral contribution.

**Figure 5.**
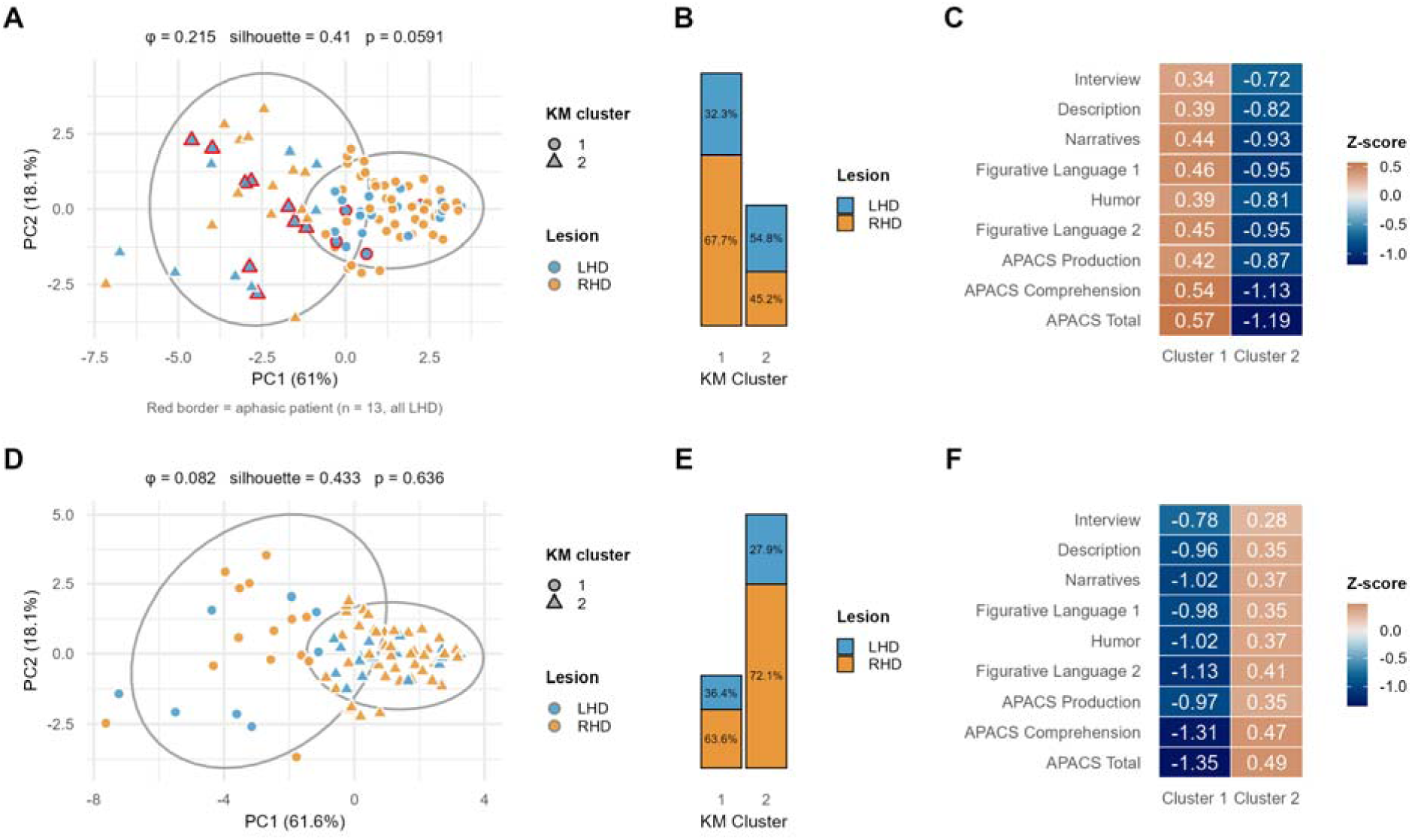
K-means clustering of stroke patients on the nine APACS measures, for the full sample (A-C) and with aphasic patients excluded (D-F). Clustering used k = 2 on z-standardized scores (available-case: full n = 96; aphasia excluded n = 83). **(A, D)** Principal-component projection; points colored by lesion side (LHD, orange; RHD, blue) and shaped by cluster, with 95% ellipses. PC1/PC2 explain 61.0%/18.1% (full) and 61.6%/18.1% (aphasia excluded). Dark magenta borders mark the 13 aphasic patients (all LHD; one excluded for missing data). **(B, E)** Within-cluster composition by lesion side. **(C, F)** Cluster-center profiles as standardized means per measure; warm = above-average, cool = below-average performance.

#### Correlations of APACS Scores with Other Neuropsychological and Cognitive Measures

Theory of Mind (SET) was a robust correlate in both groups (LHD r = 0.59; RHD r = 0.59; both p_FDR_ ≤ 0.005), whereas the other associations were lesion-side specific: structural language skills (ACE-R Language) related to pragmatic performance only in LHD (r = 0.82, p_FDR_ = 0.005), and abstract reasoning (RAVEN) only in RHD (r = 0.62, p_FDR_ = 0.003) (Figure 6A, B). Theory of Mind thus appears to be a shared substrate across hemispheres, while structural language and general reasoning differed between LHD and RHD. Robustness profiling supported this reading: the four FDR-significant associations were all rated High. However, it should be noted that the LHD ACE-R Language correlation is based almost entirely on aphasic patients (8 of the 10 LHD cases with data), so linguistic ability and aphasia status are mixed together and cannot be disentangled in this sample. Finally, in an exploratory whole-sample analysis on overall pragmatic abilities, APACS Total showed a weak positive association with self-rated communicative effectiveness (COAST Patient Total; r = 0.33, p = 0.03, N = 43; Figure 6C, D). However, its robustness was Low (bootstrap CI marginally including zero, Pearson-Spearman divergence), so it is best taken as preliminary evidence that patients’ perceived communicative effectiveness tracks overall APACS performance regardless of lesion side.

**Figure 6.**
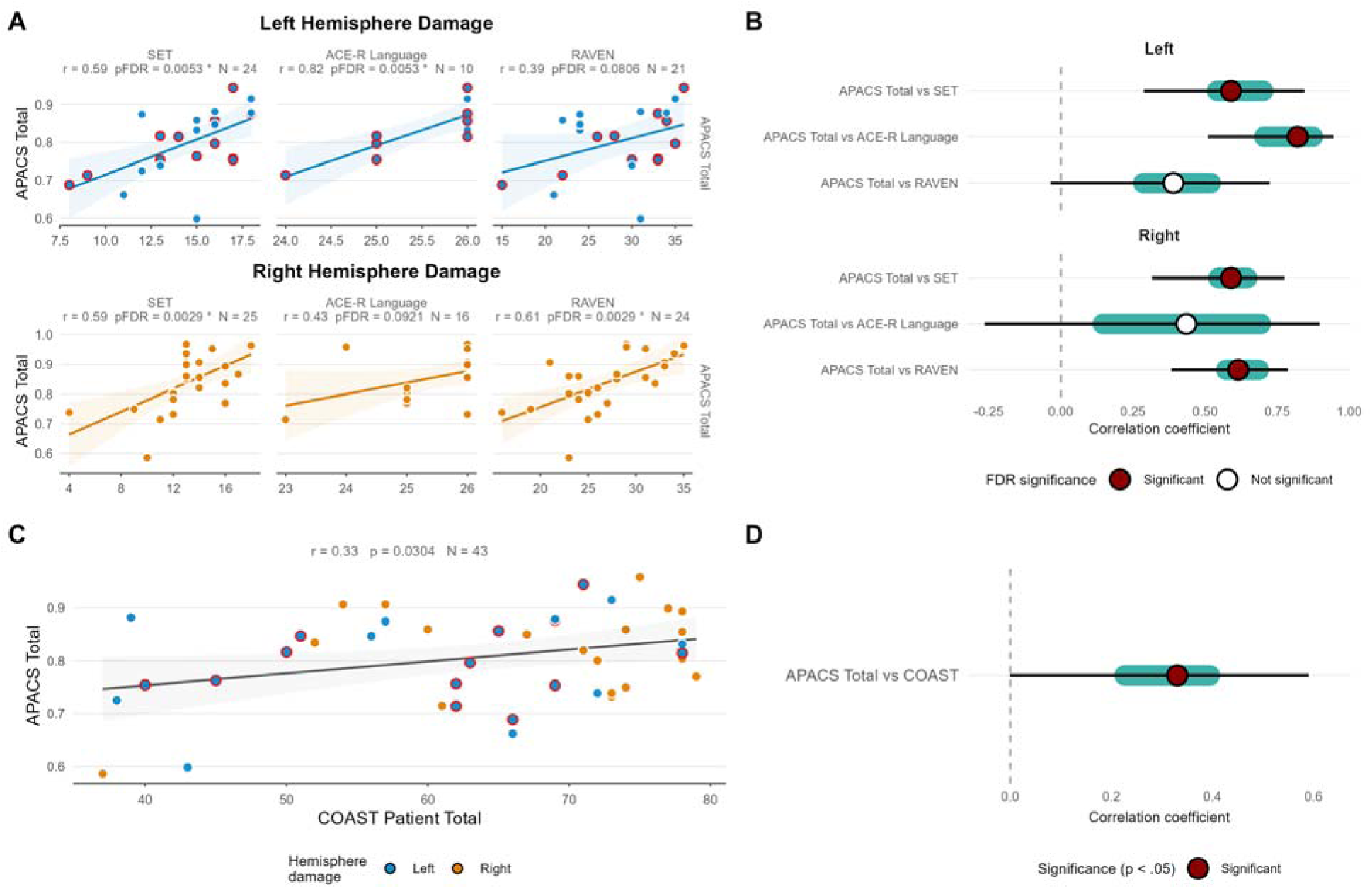
Correlations between APACS Total and cognitive/communicative measures, with robustness profiling. **(A)** Scatterplots of APACS Total against SET, ACE-R Language, and RAVEN, separately for LHD (top, blue) and RHD (bottom, orange); lines are OLS fits with 95% bands. Each strip reports Pearson r, FDR-corrected p, and available-case N, which varies because each correlation uses all participants with non-missing data. **(B)** Robustness profile for the same six associations: filled point = Pearson r (dark red = FDR-significant, white = non-significant), teal band = leave-one-out (LOO) interval, black whisker = bootstrap 95% CI (1,000 resamples). **(C)** Exploratory whole-sample association between APACS Total and COAST Patient Total, with points colored by lesion side; the annotation reports Pearson r, p, and N. **(D)** Corresponding robustness profile for the COAST association. Aphasic patients (all LHD, blue) are marked by a red outline.

## Discussion

Three findings emerged from this APACS-based comparison of pragmatic profiles after left- and right-hemisphere stroke. First, permutation testing, equivalence testing, and clustering converged on largely comparable performance across lesion sides, the only residual LHD disadvantage being on the two open-output tasks. Second, the cognitive basis was partly shared: Theory of Mind correlated with pragmatics in both groups, whereas structural language was tied to LHD and abstract reasoning to RHD. Third, excluding aphasic patients reduced but did not remove the LHD impairment, pointing to a pragmatic deficit that is not merely secondary to aphasia.

### Pragmatic profiles are largely comparable between LHD and RHD

Having established that stroke survivors performed worse than controls on most APACS measures, except the two near-ceiling multiple-choice tasks (Figurative Language 1 and Humor), we compared LHD and RHD. First, permutation tests showed no lesion-side difference on most measures; the only reliable differences, all favoring RHD, were observed in Interview, Figurative Language 2, and APACS Production. Second, and more informatively, equivalence testing found statistical equivalence on five of nine measures (Description, Narratives, Figurative Language 1, Humor, APACS Comprehension); the inconclusive outcomes were limited to the two open-output tasks (i.e., Interview and Figurative Language 2) and the Production and Total composites, whose narrow margins were exceeded because of limited precision rather than because of a true difference. Because a non-significant difference test cannot by itself show comparability, this equivalence supports the positive claim that the groups are alike. Third, unsupervised clustering grouped patients by overall severity rather than lesion side, with accuracy of lesion side grouping no better than chance. Taken together, the pragmatic performance profile does not track lesion side, consistent with a bilateral contribution.

This pattern fits the neuroimaging literature. Reyes-Aguilar et al. (2018) found that pragmatic comprehension relies on a bilateral fronto-temporal network, including the medial prefrontal cortex, which brings language and social-cognition regions together into a “pragmatic language network”. Also, fMRI studies and meta-analyses report bilateral activation across pragmatic tasks, with left-hemisphere core language regions active alongside right-hemisphere areas (Bambini et al., 2011; Bohrn et al., 2012; Rapp et al., 2012; Bischetti et al., 2024). More broadly, the search for simple left vs. right differences is at odds with the complexity of the nervous systems (Malatesta & Tommasi, 2023), which fails to emerge neatly even in classic linguistic tasks (e.g., Awana et al., 2026). Our comparable deficits after LHD and RHD provide lesion-based evidence of the same point: if pragmatic processing draws on a bilateral network, damage to either side can disrupt it.

The residual LHD disadvantage is informative rather than a problem for this view. The two affected tasks, Interview and Figurative Language 2, are the ones that require extended open verbal output. Equivalence testing classified them as inconclusive because their confidence intervals fell below zero, marking a real but limited difference. The simplest reading is that LHD patients are held back by the open-output demands of these tasks rather than by a deeper pragmatic deficit (Heine et al., 2014). In line with this, the receptive measures that involve no open output showed neither significant differences nor inconclusive equivalence, which ties the LHD disadvantage to expressive demands rather than to pragmatic ability itself. These findings match Spaccavento et al. (2024), who tested LHD and RHD pragmatics with the Italian version of the Protocol Montréal d’Évaluation de la Communication (MEC; Tavano et al., 2013) and also found no group differences, though their pilot sample was small (15 LHD, 7 RHD) and their conclusions tenuous. Our larger sample and robust analytical approach provide stronger evidence, showing that LHD and RHD performance is statistically equivalent overall, rather than simply undifferentiated. Overall, our results support the view that pragmatic ability is not confined to the right hemisphere but draws on a more bilateral network (Reyes-Aguilar et al., 2018; Bischetti, Frau & Bambini, 2024).

### Shared and distinct cognitive substrates of pragmatic performance

Although the behavioral profiles were comparable, the correlations showed a cognitive basis that was only partly shared. Theory of Mind (the non-verbal SET) was a strong correlate in both groups, of the same magnitude (r = 0.59), and one of the most stable in the robustness checks. The other correlates depended on lesion side: in LHD, performance was strongly tied to residual language (ACE-R Language) and only weakly to abstract reasoning (Raven, which did not survive correction); in RHD the pattern reversed, with a strong link to Raven but not to structural language. This double dissociation suggests that even when the behavioral outcome is the same, the cognitive resources behind it shift with the lesioned hemisphere.

This does not reduce pragmatics to the contribution of other skills. Recent resting-state fMRI suggests pragmatic processing is partly separate from both the ToM and the structural language networks (Forbes Schieche et al., 2025), and task-based fMRI shows that the core language network is not specifically engaged by mentalizing (Shain et al., 2023). Together, these findings point to pragmatic ability as a complex interface domain between language and cognition. The association with ToM that we found in both groups is consistent with accounts in which mentalizing supports pragmatic interpretation across tasks and populations (Papafragou & Grigoroglou, 2025), while recognizing that the extent of its contribution varies across phenomena and contexts (Bambini & Lecce, 2025). Our findings align with this perspective: ToM emerged as a shared correlate, whereas language and reasoning helped in ways that depended on lesion side, and the residual pragmatic deficit was not fully explained by either domain (Bosco, Tirassa, & Gabbatore, 2018; Bischetti, Frau, & Bambini, 2024).

### LHD pragmatic impairment is not reducible to aphasia

A key question is whether pragmatic impairment in LHD is primary, that is, a genuine pragmatic disruption, or secondary to the structural language deficits that define aphasia (Cummings, 2021). This is why aphasic patients have so often been excluded, and why we conducted every analysis twice, on the full sample and with aphasic patients removed. Removing aphasic LHD patients did not abolish pragmatic impairment: below-threshold performance remained common in non-aphasic LHD, at similar or higher rates than RHD on several measures. However, the residual lesion-side differences were attenuated: no measure survived correction in the permutation tests, the Interview and Figurative Language 2 effects became small, the composite confidence intervals that had previously sat just outside the margin in equivalence tests now included zero, and the weak clustering association moved further toward chance.

Two conclusions follow. First, the aphasic patients in our sample, whose residual language impairment was mostly mild, contributed disproportionately to the apparent LHD disadvantage but did not entirely explain the pragmatic deficit of the whole LHD group. This fits the clustering, which placed most of them in the lower-performing cluster, a cluster which also included non-aphasic LHD and RHD patients. Second, and more important for the primary-vs-secondary question, below-threshold pragmatic performance remained widespread in non-aphasic LHD patients, indicating that pragmatic impairment in LHD stroke is not just a by-product of structural language breakdown. These patients are free of the structural-language confound, yet their pragmatic deficits are comparable to those in RHD patients, the group in which pragmatic impairment is usually expected. This provides the clearest evidence in our data for a primary pragmatic deficit in a clinically neglected group. It also matches earlier reports that pragmatic difficulties after LHD can be worse than structural language impairment alone would predict, and may not recover at the same pace as language (Jospe et al., 2022).

### Clinical implications

The bilateral pragmatic impairments we found question the traditional clinical focus on right-hemisphere damage. Patients with strokes in either hemisphere have trouble understanding non-literal communication, yet in LHD these problems are rarely assessed or diagnosed (McDonald, 2000; Cutica, Bucciarelli & Bara, 2006; Sidtis & Yang, 2017). Their difficulties are then put down to structural language deficits alone, which misses an important part of the picture. Because pragmatic processing is bilateral, pragmatic assessment should be part of standard care for all stroke patients, whatever the lesion side; a focus on RHD alone risks missing real impairments in LHD survivors and leaving them without treatment. Importantly, routine pragmatic assessment is now a realistic goal, thanks to the availability of standardized clinical batteries, such as APACS (Arcara & Bambini, 2016) and its short variant, available also in languages other than Italian (e.g., Petit et al., 2025; Wilkens et al., 2026), together with a growing repertoire of computational approaches that quantify pragmatic aspects of connected speech through natural language processing (Meister et al., 2026), enabling scalable and increasingly objective assessment in clinical settings. Further support for the everyday relevance of pragmatic assessment comes from the exploratory link between APACS Total and COAST, which was positive regardless of lesion side. It was moderate and did not fully survive the robustness checks, so it is only a preliminary but suggestive signal that the difficulties that APACS captures are noticeable to patients, and that future work should track patients’ own ratings of communication alongside performance-based tests.

### Limitations and future directions

Several limitations apply. First, although the overall sample is fairly large, the aphasic LHD subgroup was small (n = 14) and mostly mildly impaired, so our conclusions about aphasia apply mainly to mild-to-moderate cases, not severe ones. Excluding aphasic patients also shrank the LHD group unevenly (from 40 to 26). Second, the cognitive and communicative tests were administered only to some patients, so the correlations rest on small, unequal samples (10 to 25 per lesion-side). This mostly affects the language and reasoning results: the LHD-Raven and RHD-structural language correlations came from small subsamples with limited robustness, so the double dissociation should be treated as a hypothesis to test in a properly powered study, not as a settled result. We handled this in part through available-case analysis and an explicit robustness framework that weighs robustness ratings together with significance, but replication with complete batteries is needed. The association of pragmatics with COAST is likewise preliminary. Third, the cross-sectional, post-acute design cannot address recovery over time or whether the LHD open-output disadvantage fades as language recovers. Fourth, lesion data were available for most but not all patients and were coded at the level of broad anatomical regions rather than voxel-based lesion-symptom mapping; finer analyses would tie the dissociations to specific neural substrates.

## Conclusion

The present study challenges prevailing assumptions about hemispheric contributions to pragmatics in stroke. Remarkably, pragmatic performance was comparable at the behavioral level in both groups, despite partly different underlying cognitive domains: Theory of Mind was a common factor, while LHD performance relied more on residual linguistic abilities and RHD more on abstract reasoning. Critically, pragmatic impairment in LHD persisted after excluding aphasic patients, indicating that it is not merely secondary to structural language breakdown as most often assumed, but can represent a primary deficit following left sided stroke. These findings highlight the importance of extending pragmatic assessment to all stroke patients, regardless of lateralization, to guide more targeted interventions.

## CRediT author statement

**Simone Gastaldon:** Conceptualization, Methodology, Software, Validation, Formal Analysis, Data Curation, Writing - Original Draft, Visualization. **Fortunata Romeo:** Investigation, Data Curation, Writing - Review and Editing. **Chiara Barattieri di San Pietro:** Methodology, Data Curation, Writing - Review and Editing. **Natalia Chumakova:** Investigation, Data Curation. **Daniela D’Imperio:** Investigation, Data Curation. **Sara Lago:** Investigation, Data Curation. **Sara Nordio:** Investigation, Data Curation. **Ilaria Parrotta:** Investigation, Data Curation. **Marco Rigoni:** Resources, Project Administration, Writing - Review and Editing. **Valentina Bambini:** Conceptualization, Methodology, Resources, Supervision, Writing - Review and Editing. **Giorgio Arcara:** Conceptualization, Methodology, Resources, Project Administration, Writing - Review and Editing.

## Acknowledgements

We would like to thank Laura Passarini, Martina Garzon, Francesca Meneghello, and Marco Berlot for their support in coordinating and conducting data collection.

## Data availability statement

Data are not publicly available due to privacy and data protection regulations. Access to the data is restricted because they contain sensitive information that could compromise participant confidentiality. Requests for access may be considered by the corresponding author, subject to applicable legal, ethical, and institutional requirements

## References

Angeleri, R., Bosco, F. M., Zettin, M., Sacco, K., Colle, L., & Bara, B. G. (2012). Assessment battery for communication (ABaCo): Normative data. Behavior Research Methods, 44(3), 845–861. 10.3758/s13428-011-0174-9

Arcara, G., & Bambini, V. (2016). A test for the assessment of pragmatic abilities and cognitive substrates (APACS): Normative data and psychometric properties. Frontiers in Psychology, 7, 70. 10.3389/fpsyg.2016.00070

Arcara, G., Tonini, E., Muriago, G., Mondin, E., Sgarabottolo, E., Bertagnoni, G., Semenza, C., & Bambini, V. (2020). Pragmatics and figurative language in individuals with traumatic brain injury: Fine-grained assessment and relevance-theoretic considerations. Aphasiology, 34(8), 1070– 1100. 10.1080/02687038.2019.1615033

Awana, A., Berthier, M. L., Torres-Prioris, M. J., & López-Barroso, D. (2026). Mapping the neural patterns of verbal repetition: An activation likelihood estimation meta-analysis. Brain Structure and Function, 231(5), 75. 10.1007/s00429-026-03119-3

Bambini, V., Gentili, C., Ricciardi, E., Bertinetto, P. M., & Pietrini, P. (2011). Decomposing metaphor processing at the cognitive and neural level through functional magnetic resonance imaging. Brain Research Bulletin, 86(3), 203–216. 10.1016/j.brainresbull.2011.07.015

Bambini, V., Arcara, G., Martinelli, I., Bernini, S., Alvisi, E., Moro, A., Cappa, S., & Ceroni, M. (2016a). Communication and pragmatic breakdowns in amyotrophic lateral sclerosis patients. Brain and Language, 153*–*154, 1–12. 10.1016/j.bandl.2015.12.002

Bambini, V., Arcara, G., Bechi, M., Buonocore, M., Cavallaro, R., & Bosia, M. (2016b). The communicative impairment as a core feature of schizophrenia: Frequency of pragmatic deficit, cognitive substrates, and relation with quality of life. Comprehensive Psychiatry, 71, 106–120. 10.1016/j.comppsych.2016.08.012

Bambini, V., Arcara, G., Aiachini, B., Cattani, B., Dichiarante, M. L., Moro, A., Cappa, S. F., & Pistarini, C. (2017). Assessing functional communication: Validation of the Italian versions of the Communication Outcome after Stroke (COAST) scales for speakers and caregivers. Aphasiology, 31(3), 332–358. 10.1080/02687038.2016.1225275

Bambini, V., Bianco, M., Stella, M., Bosia, M., Buonocore, M., & Cavallaro, R. (2020). A leopard cannot change its spots: A novel pragmatic account of concretism in schizophrenia. Neuropsychologia, 139, 107332. 10.1016/j.neuropsychologia.2020.107332

Bambini, V., & Lecce, S. (2025). At the heart of human communication: New views on the complex relationship between pragmatics and Theory of Mind. Philosophical Transactions of the Royal Society B: Biological Sciences, 380(1932), 20230486. 10.1098/rstb.2023.0486

Bays, C. L. (2001). Quality of life of stroke survivors: A research synthesis. Journal of Neuroscience Nursing, 33(6), 310–316. 10.1097/01376517-200112000-00005

Benjamini, Y., & Hochberg, Y. (1995). Controlling the false discovery rate: A practical and powerful approach to multiple testing. Journal of the Royal Statistical Society: Series B (Methodological), 57(1), 289–300. 10.1111/j.2517-6161.1995.tb02031.x

Bischetti, L., Frau, F., & Bambini, V. (2024). Neuropragmatics. In M. J. Ball, N. Müller, & E. Spencer (Eds.), The handbook of clinical linguistics (2nd ed., pp. 41–54). Wiley. 10.1002/9781119875949

Blake, M. L. (2017). Right-hemisphere pragmatic disorders. In L. Cummings (Ed.), Research in clinical pragmatics (pp. 243–266). Springer. 10.1007/978-3-319-47489-2_10

Blake, M. L., Ferré, P., Murray, L., Hewetson, R., Johnson, M., Durfee, A. Z., Minga, J., Sheppard, S. M., & Love, A. (2026). Understanding right-hemisphere language disorders through a half-century of research: A systematic review. American Journal of Speech-Language Pathology. 10.1044/2026_AJSLP-25-00294

Bohrn, I. C., Altmann, U., & Jacobs, A. M. (2012). Looking at the brains behind figurative language: A quantitative meta-analysis of neuroimaging studies on metaphor, idiom, and irony processing. Neuropsychologia, 50(11), 2669–2683. 10.1016/j.neuropsychologia.2012.07.021

Borod, J. C., Rorie, K. D., Pick, L. H., Bloom, R. L., Andelman, F., Campbell, A. L., & Sliwinski, M. (2000). Verbal pragmatics following unilateral stroke: Emotional content and valence. Neuropsychology, 14(1), 112–119. 10.1037//0894-4105.14.1.112

Bosco, F. M., Tirassa, M., & Gabbatore, I. (2018). Why pragmatics and theory of mind do not (completely) overlap. Frontiers in Psychology, 9, 1453. 10.3389/fpsyg.2018.01453

Brownell, H., Michel, D., Powelson, J. A., & Gardner, H. (1983). Surprise but not coherence: Sensitivity to verbal humor in right-hemisphere patients. Brain and Language, 18(1), 20–27. 10.1016/0093-934x(83)90002-0

Caldwell, A. R. (2022). Exploring equivalence testing with the updated TOSTER R package. PsyArXiv. 10.31234/osf.io/ty8de

Campbell, G. B., Skidmore, E. R., Whyte, E. M., & Matthews, J. T. (2015). Overcoming practical challenges to conducting clinical research in the inpatient stroke rehabilitation setting. Topics in Stroke Rehabilitation, 22(5), 386–394. 10.1179/1074935714Z.0000000045

Cappelli, G., Noccetti, S., Arcara, G., & Bambini, V. (2018). Pragmatic competence and its relationship with the linguistic and cognitive profile of young adults with dyslexia. Dyslexia, 24(3), 294–306. 10.1002/dys.1588

Carlesimo, G. A., Caltagirone, C., Gainotti, G., Fadda, L., Gallassi, R., Lorusso, S., et al. (1996). The mental deterioration battery: Normative data, diagnostic reliability and qualitative analysis of cognitive impairment. European Neurology, 36(6), 378–384. 10.1159/000117297

Carotenuto, A., Arcara, G., Orefice, G., Cerillo, I., Giannino, V., Rasulo, M., Iodice, R., & Bambini, V. (2018). Communication in multiple sclerosis: Pragmatic deficit and its relation with cognition and social cognition. Archives of Clinical Neuropsychology, 33(2), 194–205. 10.1093/arclin/acx061

Catani, M., & Bambini, V. (2014). A model for social communication and language evolution and development (SCALED). Current Opinion in Neurobiology, 28, 165–171. 10.1016/j.conb.2014.07.018

Champagne-Lavau, M., Stip, E., & Joanette, Y. (2007). Language functions in right-hemisphere damage and schizophrenia: Apparently similar pragmatic deficits may hide profound differences. Brain, 130, e67. 10.1093/brain/awl311

Champagne-Lavau, M., & Joanette, Y. (2009). Pragmatics, theory of mind and executive functions after a right-hemisphere lesion: Different patterns of deficits. Journal of Neurolinguistics, 22(5), 413–426. 10.1016/j.jneuroling.2009.02.002

Chiappe, D. L., & Chiappe, P. (2007). The role of working memory in metaphor production and comprehension. Journal of Memory and Language, 56(2), 172–188. 10.1016/j.jml.2006.11.006

Cliff, N. (1993). Dominance statistics: Ordinal analyses to answer ordinal questions. Psychological Bulletin, 114(3), 494–509. 10.1037/0033-2909.114.3.494

Coelho, C., & Flewellyn, L. (2003). Longitudinal assessment of coherence in an adult with fluent aphasia: A follow-up study. Aphasiology, 17(2), 173–182. 10.1080/729255216

Crawford, J. R., & Garthwaite, P. H. (2006). Comparing patients’ predicted test scores from a regression equation with their obtained scores: A significance test and point estimate of abnormality with accompanying confidence limits. Neuropsychology, 20(3), 259–271. 10.1037/0894-4105.20.3.259

Cummings, L. (2015). Theory of mind in utterance interpretation: The case from clinical pragmatics. Frontiers in Psychology, 6, 1286. 10.3389/fpsyg.2015.01286

Cummings, L. (Ed.). (2021). Handbook of pragmatic language disorders: Complex and underserved populations. Springer. 10.1007/978-3-030-74985-9

Cutica, I., Bucciarelli, M., & Bara, B. G. (2006). Neuropragmatics: Extralinguistic pragmatic ability is better preserved in left-hemisphere-damaged patients than in right-hemisphere-damaged patients. Brain and Language, 98(1), 12–25. 10.1016/j.bandl.2006.01.001

Deighton, S., Ju, N., Graham, S. A., Yeats, K. O. (2020). Pragmatic language comprehension after pediatric traumatic brain injury: A scoping review. The Journal of Head Trauma Rehabilitation, 35(2), E113–E126. 10.1097/HTR.0000000000000515

Dodich, A., Cerami, C., Canessa, N., Crespi, C., Iannaccone, S., Marcone, A., et al. (2015). A novel task assessing intention and emotion attribution: Italian standardization and normative data of the story-based empathy task. Neurological Sciences, 36(10), 1907–1912. 10.1007/s10072-015-2281-3

Ferré, P., Fonseca, R. P., Ska, B., & Joanette, Y. (2012). Communicative clusters after a right-hemisphere stroke: Are there universal clinical profiles? Folia Phoniatrica et Logopaedica, 64(4), 199–207. 10.1159/000340017

Ferstl, E. C., Neumann, J., Bogler, C., & von Cramon, D. Y. (2008). The extended language network: A meta-analysis of neuroimaging studies on text comprehension. Human Brain Mapping, 29(5), 581–593. 10.1002/hbm.20422

Forbes Schieche, C., Mahal, M., Thompson, W. H., & Uddén, J. (2025). Pragmatics partially segregated from Theory of Mind: Evidence from resting-state functional connectivity. Philosophical Transactions of the Royal Society B: Biological Sciences, 380(1932). 10.1098/rstb.2023.0498

Frank, C. K. (2018). Reviving pragmatic theory of theory of mind. AIMS Neuroscience, 5(2), 116–131. 10.3934/Neuroscience.2018.2.116

Frau, F., Bosia, M., Bischetti, L., Cappelli, G., Carotenuto, A., Diamanti, L., Montemurro, S., Agostoni, G., Bechi, M., D’Imperio, D., Lago, S., Noccetti, S., Simi, N., Ceroni, M., Iodice, R., Signorini, M., Arcara, G., & Bambini, V. (2025). Ten years of using the APACS test: A multistudy cross-diagnostic analysis of pragmatic profiles and their relationship with Theory of Mind. Philosophical Transactions of the Royal Society B, 380, 20230495. 10.1098/rstb.2023.0495

Fridriksson, J., Nettles, C., Davis, M., Morrow, L., & Montgomery, A. (2006). Functional communication and executive function in aphasia. Clinical Linguistics & Phonetics, 20(6), 401–410. 10.1080/02699200500075781

Fridriksson, J., & Hillis, A. E. (2021). Current approaches to the treatment of post-stroke aphasia. Journal of Stroke, 23(2), 183–201. 10.5853/jos.2020.05015

Gardner, H., & Brownell, H. H. (1986). Right Hemisphere Communication Battery. Psychology Service.

García, E. L., Ferré, P., & Joanette, Y. (2021). Right-hemisphere language disorders. In L. Cummings (Ed.), Handbook of pragmatic language disorders. Springer. 10.1007/978-3-030-74985-9_12

Glosser, G., & Goodglass, H. (1990). Disorders in executive control functions among aphasic and other brain-damaged patients. Journal of Clinical and Experimental Neuropsychology, 12(4), 485–501. 10.1080/01688639008400995

Grice, H. P. (1989). Studies in the way of words. Harvard University Press.

Heine, B., Kuteva, T., & Kaltenöck, G. (2014). Discourse, grammar, the dual process model, and brain lateralization: Some correlations. Language and Cognition, 6, 146–180. 10.1017/langcog.2013.3

Herpich, F., & Rincon, F. (2020). Management of acute ischemic stroke. Critical Care Medicine, 48(11), 1654. 10.1097/CCM.0000000000004597

Joanette, Y., Goulet, P., & Hannequin, D. (1990). Right hemisphere and verbal communication. Springer-Verlag.

Jospe, K., et al. (2022). Impaired empathic accuracy following damage to the left hemisphere. Biological Psychology, 172, 108380. 10.1016/j.biopsycho.2022.108380

Karamyan, V. T. (2023). Clinically applicable experimental design and considerations for stroke recovery preclinical studies. Methods in Molecular Biology, 2616, 369–377. 10.1007/978-1-0716-2926-0_25

Kasher, A., Batori, G., Soroker, N., Graves, D., & Zaidel, E. (1999). Effects of right- and left-hemisphere damage on understanding conversational implicature. Brain and Language, 68(3), 566–590. 10.1006/brln.1999.2129

Lago, S., et al. (2022). Case report: Pragmatic impairment in multiple sclerosis after worsening of clinical symptoms. Frontiers in Psychology, 13, 1028814. 10.3389/fpsyg.2022.1028814

Lakens, D. (2017). Equivalence tests: A practical primer for t tests, correlations, and meta-analyses. Social Psychological and Personality Science, 8(4), 355–362. 10.1177/1948550617697177

Lakens, D., Scheel, A. M., & Isager, P. M. (2018). Equivalence testing for psychological research: A tutorial. Advances in Methods and Practices in Psychological Science, 1(2), 259–269. 10.1177/2515245918770963

Lezak, M. D., Howieson, D. B., Bigler, E. D., & Tranel, D. (2012). Neuropsychological assessment (5th ed.). Oxford University Press.

Malatesta, G., & Tommasi, L. (2023). Editorial: Expert opinion in environmental and genetic factors impacting functional brain lateralization in development and evolution. Frontiers in Behavioral Neuroscience, 17. 10.3389/fnbeh.2023.1215176

Mar, R. A. (2004). The neuropsychology of narrative: Story comprehension, story production and their interrelation. Neuropsychologia, 42(10), 1414–1434. 10.1016/j.neuropsychologia.2003.12.016

Martin, I., & McDonald, S. (2003). Weak coherence, no theory of mind, or executive dysfunction? Solving the puzzle of pragmatic language disorders. Brain and Language, 85(3), 451–466. 10.1016/s0093-934x(03)00070-1

McDonald, S. (2000). Exploring the cognitive basis of right-hemisphere pragmatic language disorders. Brain and Language, 75(1), 82–107. 10.1006/brln.2000.2342

Meissel, K., & Yao, E. S. (2024). Using Cliff’s Delta as a non-parametric effect size measure: An accessible web app and R tutorial. *Practical Assessment*, Research, and Evaluation, 29(1). 10.7275/pare.1977

Meister, F., Sellier Silva, M., Melshin, G., El Mouslih, C., Zaher, F., Sattari, R., Wei, H. T., Mekideche, N., Bambini, V., Voppel, A., & Palaniyappan, L. (2026). Expressive pragmatic language in mood and psychotic disorders: A systematic review and meta-analysis. Schizophrenia, 12(1), 31. 10.1038/s41537-026-00733-2

Mioshi, E., Dawson, K., Mitchell, J., Arnold, R., & Hodges, J. R. (2006). The Addenbrooke’s Cognitive Examination Revised (ACE-R): A brief cognitive test battery for dementia screening. International Journal of Geriatric Psychiatry, 21(11), 1078–1085. 10.1002/gps.1610

Mondini, S., Cappelletti, M., & Arcara, G. (2022). Methodology in neuropsychological assessment: An interpretative approach to guide clinical practice. Routledge. 10.4324/9781003195221

Montemurro, S., Mondini, S., Signorini, M., Marchetto, A., Bambini, V., & Arcara, G. (2019). Pragmatic language disorder in Parkinson’s disease and the potential effect of cognitive reserve. Frontiers in Psychology, 10, 1220. 10.3389/fpsyg.2019.01220

Nordio, S., Zampieri, M., Bambini, V., & Arcara, G. (2025). Language in multiple sclerosis. In L. Cummings (Ed.), Oxford handbook of communication disorders in neurodegenerative diseases. Oxford University Press. 10.1093/oxfordhb/9780198888482.013.0015

Parola, A., Gabbatore, I., Bosco, F. M., Bara, B. G., Cossa, F. M., Gindri, P., & Sacco, K. (2016). Assessment of pragmatic impairment in right hemisphere damage. Journal of Neurolinguistics, 39, 10–25. 10.1016/j.jneuroling.2015.12.003

Papafragou, A., & Grigoroglou, M. (2025). Pragmatic communication and Theory of Mind. Philosophical Transactions of the Royal Society B: Biological Sciences, 380(1932), 20230502. 10.1098/rstb.2023.0502

Paunov, A. M., Blank, I. A., Jouravlev, O., Mineroff, Z., Gallée, J., & Fedorenko, E. (2022). Differential tracking of linguistic vs. mental state content in naturalistic stimuli by language and Theory of Mind (ToM) brain networks. Neurobiology of Language, 3(3), 413–440. 10.1162/nol_a_00071

Pessoa, L. (2022). The entangled brain: How perception, cognition, and emotion are woven together. The MIT Press. 10.7551/mitpress/14636.001.0001

Petit, N., Mengarelli, F., Geoffray Cassar, M.-M., Arcara, G., & Bambini, V. (2025). When Do Pragmatic Abilities Peak? Assessment of Pragmatic Abilities and Cognitive Substrates–French Version Psychometric Properties Across the Lifespan. Journal of Speech, Language, and Hearing Research, 68(11), 5493–5504. 10.1044/2025_JSLHR-24-00844

Premack, D., & Woodruff, G. (1978). Does the chimpanzee have a theory of mind? Behavioral and Brain Sciences, 1(4), 515–526. 10.1017/S0140525X00076512

R Core Team. (2023). R: A language and environment for statistical computing. R Foundation for Statistical Computing. https://www.R-project.org/

Rapp, A. M., Mutschler, D. E., & Erb, M. (2012). Where in the brain is nonliteral language? A coordinate-based meta-analysis of functional MRI studies. NeuroImage, 63(1), 600–610. 10.1016/j.neuroimage.2012.06.022

Reyes-Aguilar, A., Valles-Capetillo, E., & Giordano, M. (2018). A quantitative meta-analysis of neuroimaging studies of pragmatic language comprehension: In search of a universal neural substrate. Neuroscience, 395, 60–88. 10.1016/j.neuroscience.2018.10.043

Roca, M., et al. (2012). The relationship between executive functions and fluid intelligence in Parkinson’s disease. Psychological Medicine, 42(11), 2445–2452. 10.1017/S0033291712000451

Rousseaux, M., Daveluy, W., & Kozlowski, O. (2010). Communication in conversation in stroke patients. Journal of Neurology, 257(7), 1099–1107. 10.1007/s00415-010-5469-8

Rowley, D. A., Rogish, M., Alexander, T., & Riggs, K. J. (2017). Cognitive correlates of pragmatic language comprehension in adult traumatic brain injury: A systematic review and meta-analyses. Brain Injury, 31(12), 1564–1574. 10.1080/02699052.2017.1341645

Searle, J. R. (1979). Expression and meaning: Studies in the theory of speech acts. Cambridge University Press. 10.1017/cbo9780511609213

Shain, C., Paunov, A., Chen, X., Lipkin, B., & Fedorenko, E. (2023). No evidence of theory of mind reasoning in the human language network. Cerebral Cortex, 33(10), 6299–6319. 10.1093/cercor/bhac505

Sheppard, S. M., & Sebastian, R. (2021). Diagnosing and managing post-stroke aphasia. Expert Review of Neurotherapeutics, 21(2), 221–234. 10.1080/14737175.2020.1855976

Sidtis, D. V. L., & Yang, S. Y. (2017). Formulaic language performance in left- and right-hemisphere damaged patients: Structured testing. Aphasiology, 31(1), 82–99. 10.1080/02687038.2016.1157136

Spaccavento, S., Caliendo, S., Galetta, R., Picciola, E., Losavio, E., & Glueckauf, R. (2024). Pragmatic communication deficit and functional outcome in patients with right- and left-brain damage: A pilot study. Brain Sciences, 14(4), 387. 10.3390/brainsci14040387

Sperber, D., & Wilson, D. (1995). Relevance: Communication and cognition (2nd ed.). Blackwell.

Sperber, D., & Wilson, D. (2002). Pragmatics, modularity and mind-reading. Mind & Language, 17(1–2), 3–23. 10.1111/1468-0017.00186

Tavano, A., Côté, H., Ferré, P., Ska, B., & Joanette, Y. (2013). Protocollo MEC: Protocollo Montréal per la valutazione delle abilità comunicative. Springer. 10.1007/978-88-470-5456-1

Tomasello, R., Boux, I., & Pulvermüller, F. (2025). Theory of Mind and the brain substrates of direct and indirect communicative action understanding. Philosophical Transactions of the Royal Society B: Biological Sciences, 380(1932), 20230497. 10.1098/rstb.2023.0497

Tsolakopoulos, D., Kasselimis, D., Laskaris, N., Angelopoulou, G., Papageorgiou, G., Velonakis, G., Varkanitsa, M., Tountopoulou, A., Vassilopoulou, S., Goutsos, D., & Potagas, C. (2023). Exploring pragmatic deficits in relation to theory of mind and executive functions: Evidence from individuals with right hemisphere stroke. Brain Sciences, 13(10), 1385. 10.3390/brainsci13101385

Vargha, A., & Delaney, H. D. (2000). A critique and improvement of the CL common language effect size statistics of McGraw and Wong. Journal of Educational and Behavioral Statistics, 25(2), 101–132. 10.3102/10769986025002101

Wilkens, L. S., Ciaccio, L. A., Picht, T., Bosia, M., Arcara, G., Bambini, V., & Tomasello, R. (2026). A rapid assessment of pragmatic abilities and cognitive substrates: The German APACS Brief. Aphasiology. [accepted manuscript]

Winner, E., & Gardner, H. (1977). The comprehension of metaphor in brain-damaged patients. Brain, 100(4), 717–729. 10.1093/brain/100.4.717

Zaidel, E., Kasher, A., Soroker, N., & Batori, G. (2002). Effects of right and left hemisphere damage on performance of the ‘Right Hemisphere Communication Battery’. Brain and Language, 80(3), 510–535. 10.1006/brln.2001.2612

