## SUPPLEMENTARY MATERIAL for "Challenging the right-hemisphere assumption in post-stroke pragmatics: largely comparable impairment profiles across lesion sides"

1. Supplementary Figures 1-3
2. Supplementary Tables 1-10

### SUPPLEMENTARY FIGURES

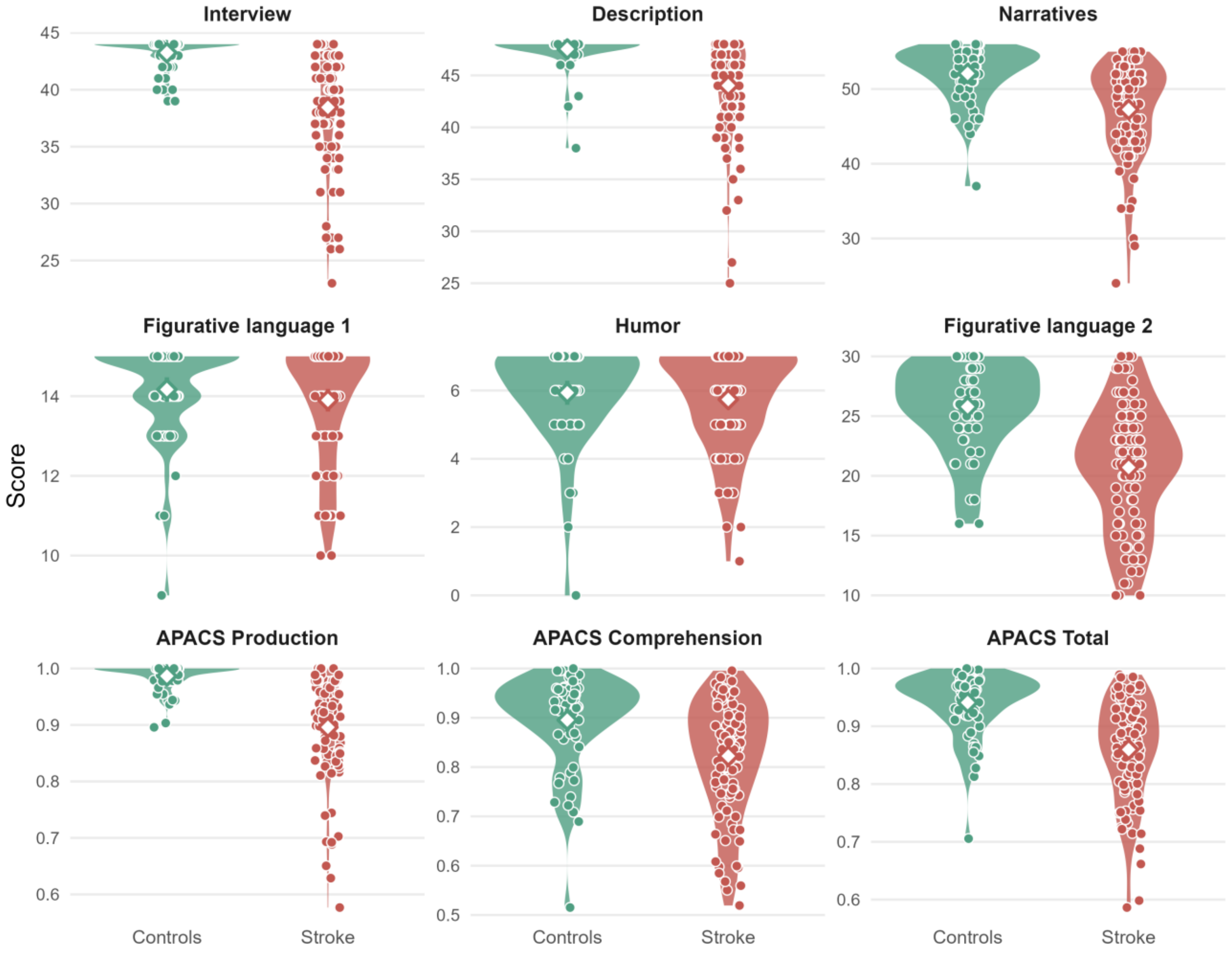


**Supplementary Figure 1.** APACS score distributions in stroke patients and healthy controls. For each task and composite score, individual scores are shown as dots, and the group mean with its 95% Cis (mean ± 1.96·SE) is marked by a white diamond with a vertical bar, over the full score distribution (violin) of each group (controls, green; stroke, red).

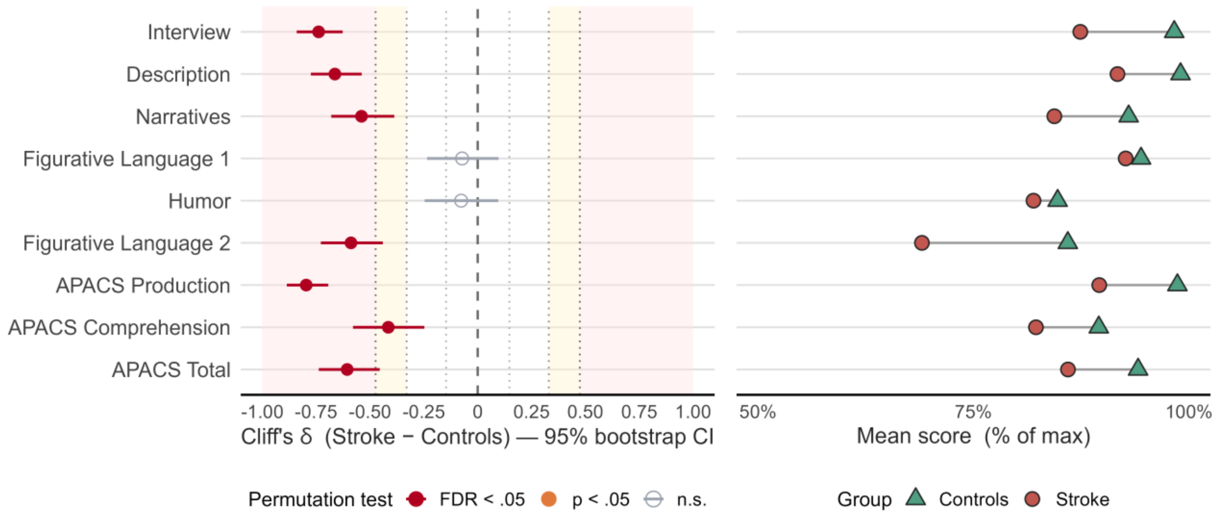


**Supplementary Figure 2.** Effect sizes of pragmatic performance in stroke patients versus healthy controls across the nine APACS measures. **(A)** Cliff's δ effect sizes with 95% bootstrap confidence intervals (5,000 resamples) for the stroke–control contrast; negative values indicate lower scores in stroke patients. Point color encodes the permutation-test outcome (20,000 resamples): filled red = significant after Benjamini-Hochberg FDR correction across the nine measures, filled orange = significant before correction only, open grey = non-significant. Background shading marks effect-size benchmarks (Vargha & Delaney, 2000: yellow = medium, $|\delta| \geq .33$; red = large, $|\delta| \geq .47$); dotted lines mark the small, medium, and large thresholds (±.15, ±.33, ±.47). Measures are ordered by effect-size magnitude. **(B)** Group mean scores expressed as percentage of the maximum, for stroke patients (red circles) and controls (green triangles). Analyses use available-case data (stroke n = 96-99; controls n = 60). Five out of seven tasks and 2 out of 3 composite scores showed significantly lower performance in stroke patients (all $p_{FDR} < 0.001$), with effects ranging from medium to large (Cliff's δ from -0.42 to -0.80); only Figurative Language 1 ($\delta = -0.07$, $p_{FDR} = 0.28$) and Humor ($\delta = -0.08$, $p_{FDR} = 0.42$) did not differ between groups.

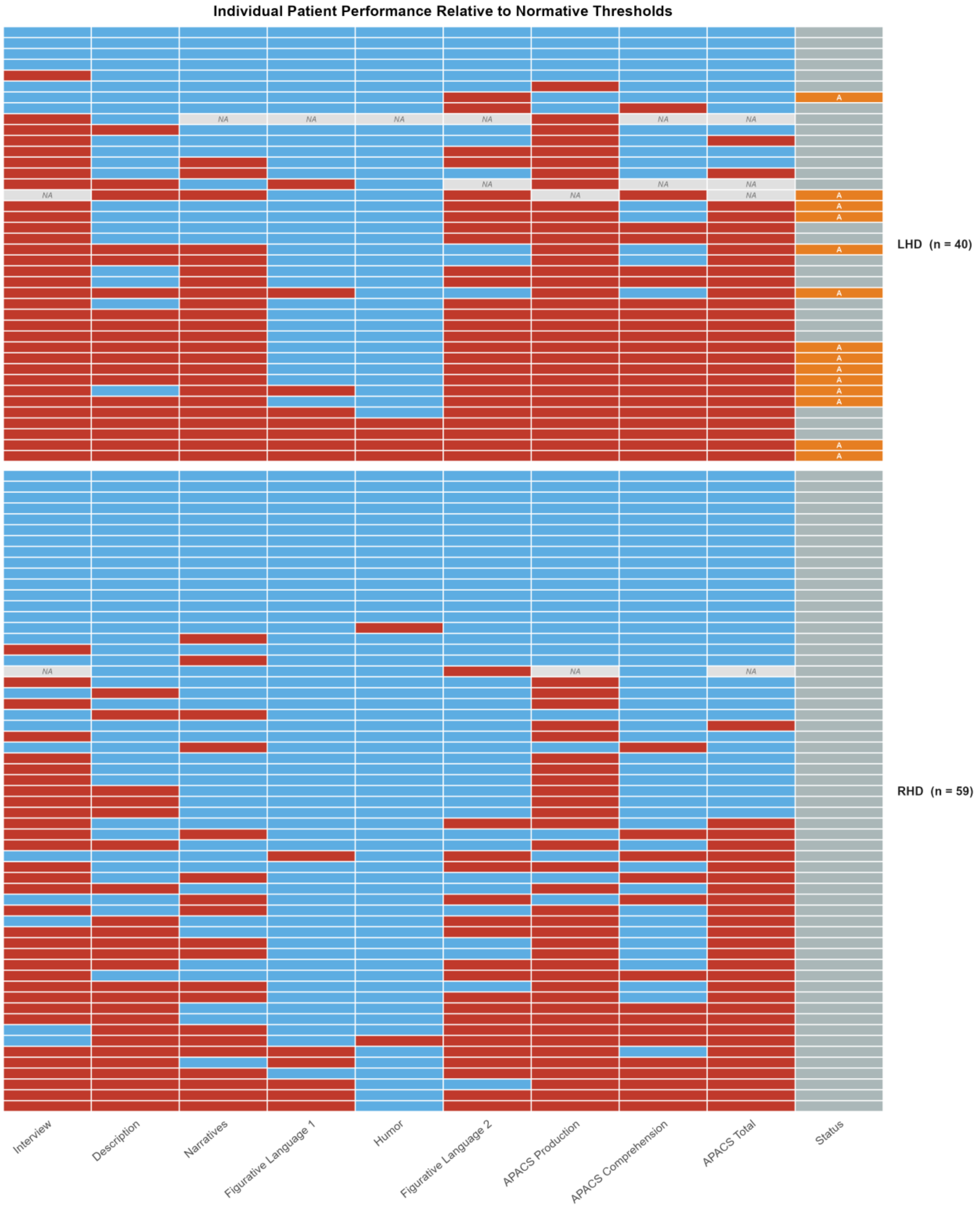


**Supplementary Figure 3.** Individual patient performance on the nine APACS measures relative to normative cut-offs. Each row represents one stroke patient and each column one APACS measure; tiles are coloured red when the patient scored below the published normative cut-off (5th percentile; Arcara & Bambini, 2016) and blue when within the normal range. Grey tiles labelled "NA" indicate measures for which a below-threshold classification could not be computed because the score or the applicable normative cut-off was unavailable. Patients are grouped by lesion side (top: LHD, n = 40; bottom: RHD, n = 59) and, within each group, ordered by the number of below-threshold measures (most to least impaired, top to bottom). The right-hand "Status" strip flags aphasic patients ("A", orange); all aphasic patients had left-hemisphere damage. LHD = left-hemisphere damage; RHD = right-hemisphere damage.

### SUPPLEMENTARY TABLES

**Supplementary Table 1.** Group comparisons (permutation tests) between stroke patients and healthy controls: per-measure means, mean difference, Cliff's δ, permutation p, and FDR-corrected p. Available-case analysis; FDR by Benjamini-Hochberg across the nine measures. Mean difference is Stroke minus Controls (composites on a 0-1 scale); Cliff's δ with 95% bootstrap CI (5,000 resamples); negative = stroke lower.

| Measure | Group means (M ± SD) | Mean difference | Cliff's δ [95% CI] | Perm p | $p_{FDR}$ |
|---|---|---|---|---|---|
| Interview | Stroke 38.45 ± 4.79<br>Controls 43.25 ± 1.35 | -4.80 | -0.74 [-0.84, -0.62] | <0.001 | <0.001 |
| Description | Stroke 44.02 ± 4.45<br>Controls 47.53 ± 1.64 | -3.51 | -0.66 [-0.78, -0.54] | <0.001 | <0.001 |
| Narratives | Stroke 47.25 ± 6.25<br>Controls 52.08 ± 3.72 | -4.83 | -0.54 [-0.68, -0.38] | <0.001 | <0.001 |
| Figurative Language 1 | Stroke 13.90 ± 1.44<br>Controls 14.17 ± 1.21 | -0.27 | -0.07 [-0.24, +0.10] | .254 | .286 |
| Humor | Stroke 5.74 ± 1.43<br>Controls 5.93 ± 1.39 | -0.20 | -0.08 [-0.24, +0.10] | .423 | .423 |
| Figurative Language 2 | Stroke 20.70 ± 5.09<br>Controls 25.78 ± 3.71 | -5.08 | -0.59 [-0.73, -0.44] | <0.001 | <0.001 |
| APACS Production | Stroke 0.896 ± 0.089<br>Controls 0.987 ± 0.024 | -0.091 | -0.80 [-0.89, -0.70] | <0.001 | <0.001 |
| APACS Comprehension | Stroke 0.822 ± 0.116<br>Controls 0.895 ± 0.094 | -0.073 | -0.42 [-0.58, -0.25] | <0.001 | <0.001 |
| APACS Total | Stroke 0.860 ± 0.088<br>Controls 0.941 ± 0.055 | -0.081 | -0.61 [-0.74, -0.46] | <0.001 | <0.001 |

**Supplementary Table 2.** Correlations between APACS Total and cognitive measures (by lesion side) and COAST (whole sample), with robustness diagnostics. Bootstrap 95% CI = 1,000 resamples; LOO = leave-one-out r range. † marks a bootstrap CI that crosses zero or an unstable LOO range. Robustness score (0-4 composite) counts: bootstrap CI excludes zero, LOO stable, Pearson-Spearman agreement, and absence of influential cases (max Cook's D within threshold). The correlation in the whole sample with COAST is a separate exploratory analysis and is not FDR-corrected. The bootstrap CI marginally includes zero and the Pearson-Spearman gap is modest; together with a single influential case this yields a Low robustness score.

| Hemisphere | Measure | N | r | p | p(FDR) | Spearman ρ | Bootstrap 95% CI | LOO r range | Max Cook's D | Infl. cases | Robustness |
|---|---|---|---|---|---|---|---|---|---|---|---|
| Left | SET | 24 | 0.590 | 0.002 | 0.005 | 0.645 | [0.29, 0.84] | [0.54, 0.70] | 0.20 | 1 | 3 (High) |
| Left | ACE-R Language | 10 | 0.821 | 0.004 | 0.005 | 0.811 | [0.51, 0.95] | [0.71, 0.87] | 0.32 | 0 | 4 (High) |
| Left | RAVEN | 21 | 0.390 | 0.081 | 0.081 | 0.476 | [-0.03, 0.72] † | [0.29, 0.52] | 0.21 | 1 | 2 (Moderate) |
| Right | SET | 25 | 0.590 | 0.002 | 0.003 | 0.611 | [0.32, 0.77] | [0.55, 0.64] | 0.82 | 2 | 3 (High) |

| | | | | | | | | | | | |
|---|---|---|---|---|---|---|---|---|---|---|---|
| Right | ACE-R Language | 16 | 0.435 | 0.092 | 0.092 | 0.440 | [-0.26, 0.90] † | [0.15, 0.69] † | 0.94 | 2 | 1 (Low) |
| Right | RAVEN | 24 | 0.615 | 0.001 | 0.003 | 0.634 | [0.38, 0.79] | [0.57, 0.69] | 0.29 | 2 | 3 (High) |
| Whole sample | COAST Patient Total | 43 (24 L/ 19 R) | 0.331 | 0.03 | - | 0.222 | [0, 0.59] | [0.23, 0.40] | 0.34 | 3 | 1 (Low) |

**Supplementary Table 3.** Permutation difference tests (Cliff's delta), LHD vs RHD - full stroke sample. Rows ordered by permutation p-value. n_LHD/n_RHD: available-case sizes. mean_diff: observed LHD-RHD difference; perm_p: two-sided permutation p (20,000 resamples); perm_p_fdr: Benjamini-Hochberg corrected p across the nine measures (primary criterion). cliffs_delta: effect size (positive = LHD higher) with 95% bootstrap CI [delta_ci_lo, delta_ci_hi]; delta_size by Vargha & Delaney (2000). A difference claim requires a significant permutation result; non-significance is not equivalence.

| Label | n_LHD | n_RHD | mean_LHD | sd_LHD | median_LHD | mean_RHD | sd_RHD | median_RHD | mean_LHD_norm | mean_RHD_norm | mean_diff | perm_p | cliffs_delta | delta_ci_lo | delta_ci_hi | perm_p_fdr | delta_size |
|---|---|---|---|---|---|---|---|---|---|---|---|---|---|---|---|---|---|
| Figurative Language 2 | 39 | 59 | 18.359 | 5.363 | 18.000 | 22.254 | 4.277 | 22.000 | 0.612 | 0.742 | -3.895 | 0.000 | -0.431 | -0.640 | -0.209 | 0.000 | medium |
| Interview | 39 | 58 | 36.282 | 5.351 | 37.000 | 39.914 | 3.752 | 41.000 | 0.825 | 0.907 | -3.632 | 0.000 | -0.428 | -0.631 | -0.210 | 0.001 | medium |
| APACS Production | 39 | 58 | 0.869 | 0.095 | 0.877 | 0.914 | 0.080 | 0.935 | 0.869 | 0.914 | -0.045 | 0.012 | -0.302 | -0.518 | -0.076 | 0.037 | small |
| APACS Total | 38 | 58 | 0.835 | 0.094 | 0.847 | 0.876 | 0.080 | 0.890 | 0.835 | 0.876 | -0.040 | 0.025 | -0.274 | -0.503 | -0.044 | 0.057 | small |
| APACS Comprehension | 39 | 59 | 0.799 | 0.120 | 0.812 | 0.838 | 0.111 | 0.868 | 0.799 | 0.838 | -0.040 | 0.101 | -0.215 | -0.437 | 0.018 | 0.181 | small |
| Figurative Language 1 | 40 | 59 | 13.650 | 1.578 | 14.000 | 14.068 | 1.324 | 15.000 | 0.910 | 0.938 | -0.418 | 0.174 | -0.142 | -0.358 | 0.079 | 0.262 | negligible |
| Narratives | 40 | 59 | 46.525 | 4.804 | 47.000 | 47.746 | 7.065 | 50.000 | 0.831 | 0.853 | -1.221 | 0.343 | -0.240 | -0.455 | -0.022 | 0.441 | small |
| Description | 40 | 59 | 43.725 | 4.291 | 45.000 | 44.220 | 4.575 | 46.000 | 0.911 | 0.921 | -0.495 | 0.607 | -0.089 | -0.323 | 0.143 | 0.682 | negligible |
| Humor | 40 | 59 | 5.725 | 1.585 | 6.000 | 5.746 | 1.321 | 6.000 | 0.818 | 0.821 | -0.021 | 1.000 | 0.036 | -0.191 | 0.264 | 1.000 | negligible |

**Supplementary Table 4.** Permutation difference tests (Cliff's delta), LHD vs RHD - aphasia excluded. Columns as in the full-sample permutation table.

| Label | n_LHD | n_RHD | mean_LHD | sd_LHD | median_LHD | mean_RHD | sd_RHD | median_RHD | mean_LHD_norm | mean_RHD_norm | mean_diff | perm_p | cliffs_delta | delta_ci_lo | delta_ci_hi | perm_p_fdr | delta_size |
|---|---|---|---|---|---|---|---|---|---|---|---|---|---|---|---|---|---|
| Interview | 26 | 58 | 37.077 | 5.599 | 38.500 | 39.914 | 3.752 | 41.000 | 0.843 | 0.907 | -2.837 | 0.007 | -0.292 | -0.540 | -0.025 | 0.062 | small |
| Figurative Language 2 | 25 | 59 | 19.360 | 5.844 | 20.000 | 22.254 | 4.277 | 22.000 | 0.645 | 0.742 | -2.894 | 0.014 | -0.285 | -0.561 | 0.005 | 0.062 | small |
| APACS Production | 26 | 58 | 0.882 | 0.104 | 0.912 | 0.914 | 0.080 | 0.935 | 0.882 | 0.914 | -0.032 | 0.122 | -0.178 | -0.440 | 0.080 | 0.366 | small |
| APACS Total | 25 | 58 | 0.854 | 0.101 | 0.874 | 0.876 | 0.080 | 0.890 | 0.854 | 0.876 | -0.021 | 0.310 | -0.120 | -0.406 | 0.174 | 0.698 | negligible |

| | | | | | | | | | | | | | | | | |
|---|---|---|---|---|---|---|---|---|---|---|---|---|---|---|---|---|
| APACS Comprehension | 25 | 59 | 0.824 | 0.116 | 0.842 | 0.838 | 0.111 | 0.868 | 0.824 | 0.838 | -0.014 | 0.607 | -0.077 | -0.356 | 0.199 | 0.970 | negligible |
| Narratives | 26 | 59 | 47.269 | 4.763 | 48.500 | 47.746 | 7.065 | 50.000 | 0.844 | 0.853 | -0.477 | 0.761 | -0.175 | -0.411 | 0.079 | 0.970 | small |
| Humor | 26 | 59 | 5.846 | 1.405 | 6.000 | 5.746 | 1.321 | 6.000 | 0.835 | 0.821 | 0.100 | 0.793 | 0.062 | -0.191 | 0.306 | 0.970 | negligible |
| Figurative Language 1 | 26 | 59 | 14.000 | 1.470 | 15.000 | 14.068 | 1.324 | 15.000 | 0.933 | 0.938 | -0.068 | 0.862 | -0.012 | -0.259 | 0.231 | 0.970 | negligible |
| Description | 26 | 59 | 44.192 | 4.648 | 46.000 | 44.220 | 4.575 | 46.000 | 0.921 | 0.921 | -0.028 | 0.981 | 0.023 | -0.241 | 0.278 | 0.981 | negligible |

**Supplementary Table 5.** TOST equivalence, SEM-anchored primary margin k = 2 - full stroke sample. Epsilon = 2 x SEM (SEM from APACS reliable-change tables; Crawford & Garthwaite 2006). Mean_Diff = LHD-RHD with 90% CI [CI90_Lower, CI90_Upper]. TOST_p_lower/upper: the two one-sided p-values. TOST_Outcome = 'Equivalent' when both one-sided p < .05 (90% CI within +/-Epsilon), else 'Inconclusive'. TOST never asserts a difference.

| Label | SEM | k_SEM | Epsilon | N_Left | N_Right | Mean_Left | Mean_Right | Mean_Diff | SD_Left | SD_Right | CI90_Lower | CI90_Upper | TOST_p_lower | TOST_p_upper | TOST_Outcome |
|---|---|---|---|---|---|---|---|---|---|---|---|---|---|---|---|
| Interview | 0.553 | 2 | 1.105 | 39 | 58 | 36.282 | 39.914 | -3.632 | 5.351 | 3.752 | -5.282 | -1.982 | 0.993 | 0 | Inconclusive |
| Description | 1.312 | 2 | 2.624 | 40 | 59 | 43.725 | 44.220 | -0.495 | 4.291 | 4.575 | -1.996 | 1.006 | 0.010 | 0 | Equivalent |
| Narratives | 3.150 | 2 | 6.300 | 40 | 59 | 46.525 | 47.746 | -1.221 | 4.804 | 7.065 | -3.202 | 0.760 | 0.000 | 0 | Equivalent |
| Figurative Language 1 | 0.861 | 2 | 1.722 | 40 | 59 | 13.650 | 14.068 | -0.418 | 1.578 | 1.324 | -0.923 | 0.087 | 0.000 | 0 | Equivalent |
| Humor | 0.842 | 2 | 1.683 | 40 | 59 | 5.725 | 5.746 | -0.021 | 1.585 | 1.321 | -0.527 | 0.486 | 0.000 | 0 | Equivalent |
| Figurative Language 2 | 1.542 | 2 | 3.083 | 39 | 59 | 18.359 | 22.254 | -3.895 | 5.363 | 4.277 | -5.602 | -2.189 | 0.785 | 0 | Inconclusive |
| APACS Production | 0.014 | 2 | 0.028 | 39 | 58 | 0.869 | 0.914 | -0.045 | 0.095 | 0.080 | -0.076 | -0.014 | 0.826 | 0 | Inconclusive |
| APACS Comprehension | 0.045 | 2 | 0.091 | 39 | 59 | 0.799 | 0.838 | -0.040 | 0.120 | 0.111 | -0.080 | 0.000 | 0.018 | 0 | Equivalent |
| APACS Total | 0.025 | 2 | 0.050 | 38 | 58 | 0.835 | 0.876 | -0.040 | 0.094 | 0.080 | -0.071 | -0.009 | 0.308 | 0 | Inconclusive |

**Supplementary Table 6.** TOST equivalence, SEM-anchored margin k = 2 - aphasia excluded. Columns as in the full-sample equivalence table (Supplementary Table 5).

| Label | SEM | k_SEM | Epsilon | N_Left | N_Right | Mean_Left | Mean_Right | Mean_Diff | SD_Left | SD_Right | CI90_Lower | CI90_Upper | TOST_p_lower | TOST_p_upper | TOST_Outcome |
|---|---|---|---|---|---|---|---|---|---|---|---|---|---|---|---|
| Interview | 0.553 | 2 | 1.105 | 26 | 58 | 37.077 | 39.914 | -2.837 | 5.599 | 3.752 | -4.870 | -0.804 | 0.921 | 0.001 | Inconclusive |
| Description | 1.312 | 2 | 2.624 | 26 | 59 | 44.192 | 44.220 | -0.028 | 4.648 | 4.575 | -1.855 | 1.799 | 0.011 | 0.009 | Equivalent |
| Narratives | 3.150 | 2 | 6.300 | 26 | 59 | 47.269 | 47.746 | -0.477 | 4.763 | 7.065 | -2.662 | 1.709 | 0.000 | 0.000 | Equivalent |
| Figurative Language 1 | 0.861 | 2 | 1.722 | 26 | 59 | 14.000 | 14.068 | -0.068 | 1.470 | 1.324 | -0.632 | 0.497 | 0.000 | 0.000 | Equivalent |
| Humor | 0.842 | 2 | 1.683 | 26 | 59 | 5.846 | 5.746 | 0.100 | 1.405 | 1.321 | -0.445 | 0.646 | 0.000 | 0.000 | Equivalent |
| Figurative Language 2 | 1.542 | 2 | 3.083 | 25 | 59 | 19.360 | 22.254 | -2.894 | 5.844 | 4.277 | -5.081 | -0.707 | 0.442 | 0.000 | Inconclusive |
| APACS Production | 0.014 | 2 | 0.028 | 26 | 58 | 0.882 | 0.914 | -0.032 | 0.104 | 0.080 | -0.071 | 0.006 | 0.582 | 0.006 | Inconclusive |
| APACS Comprehension | 0.045 | 2 | 0.091 | 25 | 59 | 0.824 | 0.838 | -0.014 | 0.116 | 0.111 | -0.060 | 0.032 | 0.004 | 0.000 | Equivalent |
| APACS Total | 0.025 | 2 | 0.050 | 25 | 58 | 0.854 | 0.876 | -0.021 | 0.101 | 0.080 | -0.059 | 0.017 | 0.109 | 0.002 | Inconclusive |

**Supplementary Table 7.** K-means clustering of the nine APACS measures vs lesion side - full sample. Top block: cluster-vs-lesion correspondence (best-match accuracy, %), association (φ), chi-square p and cluster cohesion (average silhouette); higher = clearer hemisphere-based grouping. Bottom block: sampling robustness of the best-match accuracy from a subject case bootstrap (1,000 resamples) - mean, SD and 95% percentile confidence interval [CI lower, CI upper]; a lower bound near 50% indicates accuracy close to chance.

| Metric | Value |
|---|---|
| Best-match accuracy | 63.542 |
| Phi | 0.215 |
| Chi-square p | 0.059 |
| Avg silhouette | 0.410 |
| Bootstrap accuracy mean | 62.927 |
| Bootstrap accuracy sd | 4.838 |
| Bootstrap accuracy CI lower (2.5%) | 53.125 |
| Bootstrap accuracy CI upper (97.5%) | 72.917 |
| Bootstrap accuracy min | 50.000 |
| Bootstrap accuracy max | 79.167 |
| Valid resamples | 1000 |

**Supplementary Table 8.** K-means clustering of the nine APACS measures vs lesion side - aphasia excluded. Blocks and columns as in the full-sample clustering table.

| Metric | Value |
|---|---|
| Best-match accuracy | 62.651 |
| Phi | 0.082 |
| Chi-square p | 0.636 |
| Avg silhouette | 0.433 |
| Bootstrap accuracy mean | 61.941 |
| Bootstrap accuracy sd | 5.187 |
| Bootstrap accuracy CI lower (2.5%) | 51.807 |
| Bootstrap accuracy CI upper (97.5%) | 71.114 |
| Bootstrap accuracy min | 50.602 |
| Bootstrap accuracy max | 81.928 |
| Valid resamples | 1000 |
